# Heterozygous loss of ZBTB38 leads to early embryonic lethality in mice via suppressing Nanog and Sox2

**DOI:** 10.1101/2021.08.19.456970

**Authors:** Miki Nishio, Takuya Matsuura, Shunya Hibi, Shiomi Ohta, Chio Oka, Noriaki Sasai, Yasumasa Ishida, Eishou Matsuda

**Affiliations:** Functional Genomics and Medicine, Nara Institute of Science and Technology, Ikoma, 630-0192, Japan; Development Biomedical Science, Nara Institute of Science and Technology; Cosmo Bio Co., Ltd., Tokyo, 135-0016, Japan

**Keywords:** Methyl-CpG binding protein (MBP), Nanog, Sox2, blastocyst, epiblast, inner cell mass (ICM).

## Abstract

Mammalian DNA methylation is an epigenetic modification which is involved in various biological processes, including gene expression regulation. In mice, methyltransferases are responsible for DNA methylation, which are critical for early embryogenesis. However, the significance of methyl-CpG binding proteins (MBPs) that bind methylated CpG remains largely unknown. We previously demonstrated that ZBTB38/CIBZ-a zinc finger type of MBP-is required for ES cell proliferation by positively regulating Nanog expression. However, the physiological function of ZBTB38 remains unclear. In this study, we generated conditional ZBTB38 knockout mice using Cre-loxP technology. Unexpectedly, our results showed that germline loss of the ZBTB38 single allele resulted in decreased epiblast cell proliferation and increased apoptosis shortly after implantation, leading to early embryonic lethality. We found that heterozygous loss of ZBTB38 reduced the expression of Nanog, Sox2, and the genes responsible for epiblast proliferation, differentiation, and cell viability. Despite this lethal phenotype, ZBTB38 is dispensable for ES cell establishment and identity. Together, these findings indicate that ZBTB38 is essential for early embryonic development, providing new insights into the roles of MBP in implantation.

## INTRODUCTION

Mammalian pre-implantation begins with zygotes that develop into 2 and 4 cells, followed by the formation of blastocysts that contain inner cell mass (ICM) and the trophectoderm (TE). Following implantation, ICM is further segregated into the epiblast and primitive endoderm (PrE), which develop into the embryo proper and yolk sac, respectively (Kojima *et al*, 2014). Meanwhile, TE forms the extra-embryonic ectoderm (ExE) and then gives rise to the placenta. In mice, implantation occurs at embryonic day (E) 4.5 (E4.5), early gastrulation occurs at E6.5, mid gastrulation at E7.5, and organogenesis from E8.5 (Kojima *et al*, 2014; Rossant & Tam, 2009). Previous evidence indicates that transcription factors play key roles in embryogenesis: epiblasts undergo proliferation via the core pluripotent transcription factors Nanog, Sox2, and Oct4, whereas PrE and TE undergo differentiation via Gata6 and Cdx2, respectively (Rossant & Tam, 2009; Frum & Ralston, 2015; Cantone & Fisher, 2013).

Pre- and peri-implantation embryos undergo remarkable reprogramming through epigenetics, mainly including DNA methylation and histone modifications (Cantone & Fisher, 2013; Skvortsova *et al*, 2018; Ivanova *et al*, 2021; Li, 2002; Cedar & Bergman, 2009). DNA methylation is highly dynamic during mouse embryogenesis: early embryos give rise to extensive DNA demethylation from the zygote to the blastocyst, whereas their DNA methylation is globally re-established during the implantation stage at E4.5-E6.5 (Cantone & Fisher, 2013; Messerschmidt *et al*, 2014; Smith *et al*, 2012). Notably, the DNA methylation level in ICM of the blastocyst is much higher than that in TE. This difference becomes significantly apparent by E6.5, when the epiblast performs most of the DNA methylation as compared to the lower methylation state of the ExE (Eckersley-Maslin *et al*, 2018). DNA methylation is mediated by a family of conserved DNA methyltransferases (Dnmt) and MBP (Buck-Koehntop & Defossez, 2013; Suzuki & Bird, 2008). Dnmt3a and Dnmt3b are responsible for de novo DNA methylation, whereas Dnmt1 maintains methylation patterns after DNA replication (Suzuki & Bird, 2008; Greenberg & Bourc’his, 2019). In contrast, MBPs bind to the methylate CpG and generally repress gene expression, but they are also associated with gene activation (Buck-Koehntop & Defossez, 2013; Zeng & Chen, 2019). Accumulating evidence has demonstrated that DNA methylation plays key roles in mammalian development and is involved in various biological processes, including gene regulation, transposon silencing, lineage specification, genomic imprinting, and X chromosome inactivation (Zeng & Chen, 2019; Klose & Bird, 2006). In previous studies, Dnmt3a/Dnmt3b double knockout (KO) mice and Dnmt1 KO mice resulted in embryonic lethality around E9.5 (Okano *et al*, 1999; Li *et al*, 1992), indicating that both de novo DNA methylation and its maintenance are critical for early embryo viability.

MBPs contain the MBD family (Mecp2, MBD1, MBD2, and MBD4), BTB-zinc finger family (Kaiso, ZBTB4, and ZBTB38), and KRAB-zinc finger protein ZFP57 (Buck-Koehntop & Defossez, 2013; Marchal & Miotto, 2015). To date, genetic studies on the MBP single gene KO mice except ZBTB38, double and triple MBP (Mecp2, MBD2, and kaiso) gene KO mice have demonstrated that all of these are dispensable for early embryogenesis (Guy *et al*, 2001; Zhao *et al*, 2003; Hendrich *et al*, 2001; Wong *et al*, 2002; Prokhortchouk *et al*, 2006; Roussel-Gervais *et al*, 2017; Martin Caballero *et al*, 2009). Thus, extensive functional redundancy could not explain the discrepancy between these MBPs in mouse embryogenesis. In addition, ZFP57 binds to certain imprinting control regions, and ZFP57 homozygous KO (ZFP57^-/-^) led to late embryonic (E14.5∼) lethality due to defects in the maintenance of both maternal and paternal imprints (Li *et al*, 2008). So far, the significance of MBPs at the peri-implantation stage remains unclear despite Dnmts playing key roles at this stage.

We previously showed that ZBTB38/CIBZ binds to methylated DNA via zinc fingers to repress or activate transcription via the BTB and spacer domain, respectively (Sasai *et al*, 2005; Oikawa *et al*, 2011). We also demonstrated that ZBTB38 promotes ES cell proliferation, inhibits ES cell differentiation toward the mesodermal lineage, and suppresses apoptosis in murine cells (Nishii *et al*, 2012; Kotoku *et al*, 2016; Oikawa *et al*, 2008). Intriguingly, ZBTB38 loss in ES cells decreased Nanog expression and, consequently, abrogated ES cell proliferation by inhibiting the G1/S transition (Nishii *et al*, 2012). Nanog is one of the first markers to be restricted within the epiblast, followed by SOX2 and OCT4, all of which are capable of regulating self-renewal and pluripotency in ES cells and epiblast cells (Niwa, 2007). Nanog is rapidly downregulated after the blastocyst stage but is expressed again in the posterior epiblast from E6.5 onwards (Chambers *et al*, 2003; Hart *et al*, 2004), and Sox2 is localized to the prospective neuroectoderm in the anterior epiblast (Avilion *et al*, 2003).

In humans, several lines of evidence indicate that the expression level of ZBTB38 is important for its normal physiological functions, and its misregulation leads to numerous diseases. Genome-wide association studies in humans indicated that the strongest association of human height with the SNP is the ZBTB38 gene, and this SNP in the region has the most significant correlation with its expression in blood (Gudbjartsson *et al*, 2008; Tu *et al*, 2015). Moreover, expressions of ZBTB38 are associated with the development of cancers (Kote-Jarai *et al*, 2011; Jing *et al*, 2019; de Dieuleveult *et al*, 2020) and neurodegenerative disease (Mead *et al*, 2012; Brown *et al*, 2014). However, the physiological function of ZBTB38 *in vivo*, remains unknown.

In this study, we generated ZBTB38 heterozygous KO (ZBTB38^+/-^) mice using conventional and Cre-loxP-based conditional KO (cKO) approaches to reveal the physiological role of ZBTB38 in mice.

## RESULTS

### ZBTB38 expression is up-regulated during embryogenesis, and heterozygous loss of ZBTB38 results in embryonic lethality

To gain insights into the role of ZBTB38 *in vivo*, we quantified the expression levels of ZBTB38 during embryogenesis in wild-type (WT, C57BL/6J) embryos using qRT-PCR. The results showed that ZBTB38 mRNA was expressed at a detectable level at the blastocyst stage (Fig 1A), which is consistent with results of our previous study that showed high expression of ZBTB38 in ES cells (Nishii *et al*, 2012; Kotoku *et al*, 2016). In the later stages, ZBTB38 transcripts were increased between E6.5 and E7.5 by four-fold, but decreased at E8.5, and elevated again at E9.5, and the increase lasted until the neonatal stage (Fig 1A). In addition, immunofluorescence analysis using the ZBTB38 antibody showed that ZBTB38 was predominantly expressed in the ICM of blastocysts and was largely colocalized with the ICM markers—Nanog, Sox2 (Fig 1B), and Oct4 (Appendix Fig S1A). ZBTB38 staining was also weakly detected in TE. At E6.5, ZBTB38 was found to be largely expressed in the epiblast and was partially colocalized with epiblast markers of Nanog, Sox2 (Fig 1C), and Oct4 (Appendix Fig S1B). Additionally, ZBTB38 colocalized with Sox2 at ExE (Fig 1C). The characteristic expression pattern of ZBTB38 implies that it is associated with early embryonic development with unique features.

**Figure 1.**
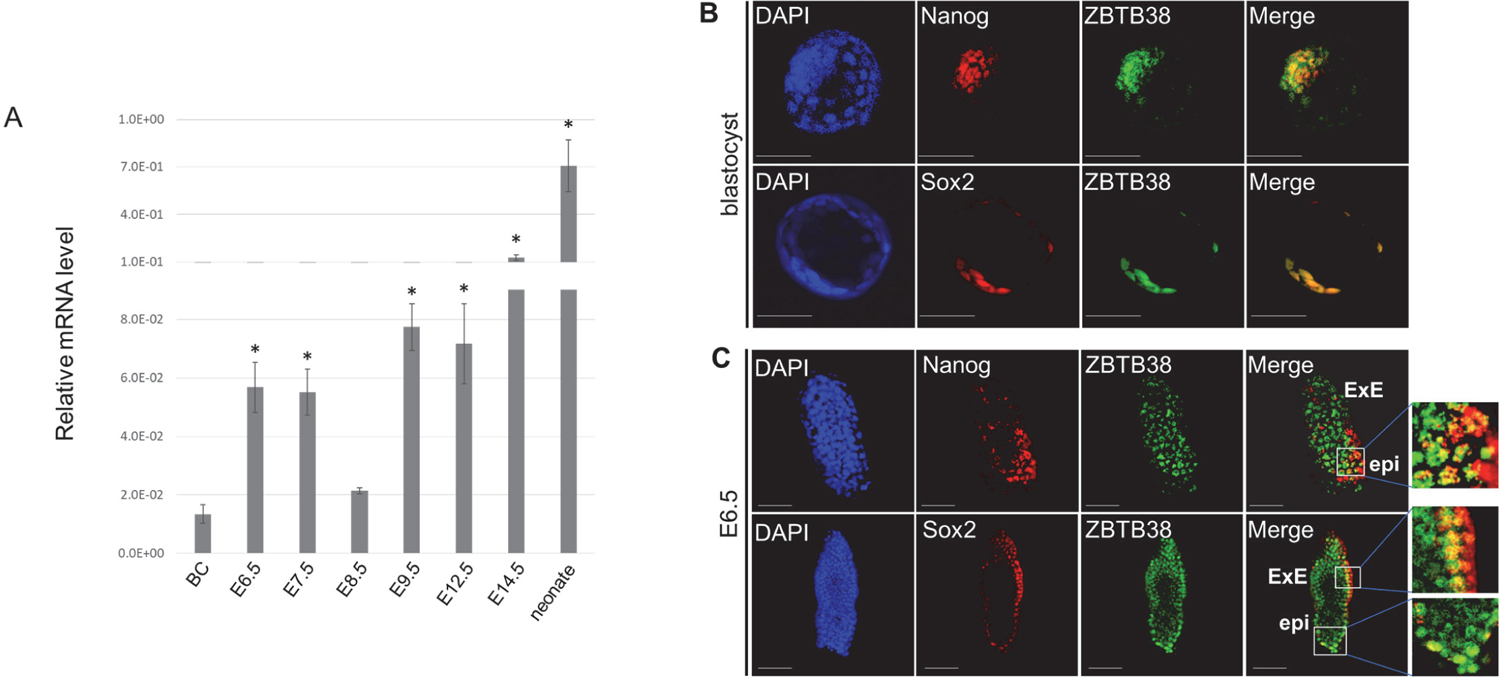
Expression of ZBTB38 during mouse embryonic development A Results of qRT-PCR for ZBTB38 expression during mouse embryonic development. Embryos from the indicated stages were carefully isolated from the uterus of C57BL/6J mice intercrosses. The TATA-binding protein (TBP) gene was used as an internal control, and the levels of transcripts were normalized against the TBP gene. BC: blastocyst. Data are representative of three independent replicates measured in triplicates, and error bars indicate ± S.D. \**p* < 0.05. B, C Whole-mount immunofluorescence and confocal microscopy for blastocyst (B) and E6.5 (C). Expressions of the indicated proteins were detected using anti-ZBTB38, anti-Nanog, and anti-Sox2 antibodies. Cell nuclei were counterstained with DAPI. Scale bar denotes 50 μm (1B) or 100 μm (1C). ExE: extraembryonic ectoderm, epi: epiblast.

To investigate the physiological role of ZBTB38 during embryogenesis, the conventional KO strategy was used to establish two independent ZBTB38^+/-^ ES cell clones that were able to maintain pluripotency and self-renewal, as described previously (Nishii *et al*, 2012). However, when the ZBTB38^+/-^ ES cells were microinjected into blastocysts to generate chimeric mice, no chimeric neonate was obtained. This finding raised the possibility that these heterozygous ES cell-derived chimera embryos did not survive in the uterus. To test this possibility and determine the timing of abnormality, we generated ZBTB38 cKO mice using the Cre-LoxP system. Two approaches were carried out to generate the ZBTB38 single null alleles (Appendix Fig S3 and the Materials and Methods). The first was crossing the ZBTB38 fl-neo/+ mice with CAG-Cre mice expressing Cre recombinase ubiquitously under the CAG promoter (Matsumura *et al*, 2004). This intercross resulted in the removal of the ZBTB38 exon3, which encodes a single transcript that produces the entire ZBTB38 protein, leading to the germline deletion of the ZBTB38 single allele (Δf-neo, Appendix Fig S4). To rule out the possibility that the remaining lacZ gene has an unexpected effect, the second approach, which requires two steps, was also performed: (A) The ZBTB38 fl-neo/+ mice were crossed with FLP transgenic mice (Kranz *et al*, 2010), leading to deletion of the lacZ and neomycin cassettes (fl, Fig 2A). (B) The ZBTB38 fl/+ mice were subsequently crossed with CAG-Cre mice, resulting in the germline deletion of the ZBTB38 single allele (Δfl, Fig 2A). Genomic PCR (gPCR) analysis revealed successful deletion of the ZBTB38 exon3 and corresponding cassettes in both methods (Fig 2B, Appendix Fig S4B), indicating that the loxP and FRT sites are functional *in vivo*.

**Figure 2.**
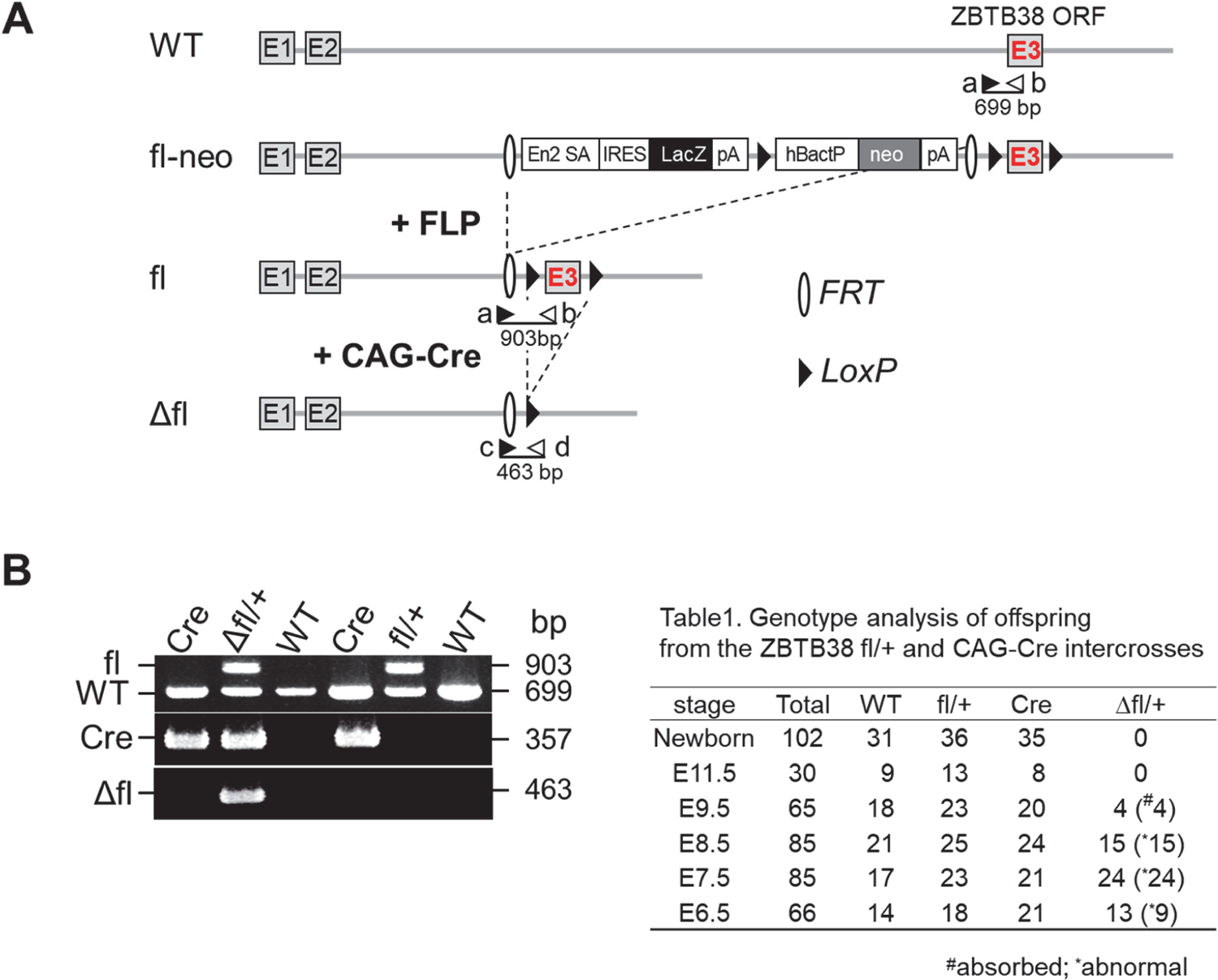
Generation of the ZBTB38 heterozygous KO mice A Schematic diagram of gene targeting strategy. Crossing the fl-neo allele to a mouse line expressing FLP recombinase removes the lacZ and neo cassettes, resulting in a conditional-ready allele (fl). When the fl allele is crossed with a mouse strain expressing CAG-Cre recombinase, exon 3 is deleted, resulting in a null allele (Δfl). Exons are shown as empty boxes and marked by a number inside. LoxP sites (black triangles) and FRT sites (empty semicircles) are shown. FLP: flp recombinase, neo: neomycine-resistant gene, En2 SA: mouse En2 splicing acceptor, IRES: internal ribosome entry site, hBactP: human b-actin promoter, pA: poly(A) signal. The position of the primers (a-d) used for genotyping is shown with arrows, and the expected sizes (bp) of genomic PCR are shown under the individual primer pairs. B PCR genotyping of E7.5 embryos isolated from the intercrosses of ZBTB38 fl/+ mice and the CAG-Cre mice. Representative PCR genotyping with primers a∼d, WT, fl, and Δfl alleles produced 699-bp, 903-bp, and 463-bp bands, respectively. The Cre genotyping primer produces a 357 bp band.

We found that either the ZBTB38 fl-neo/+ or the ZBTB38 fl/+ mice were viable, fertile, and morphologically indistinguishable from their WT or CAG-Cre littermates for at least 18 months of breeding (data not shown). Crossing the ZBTB38 fl/+ with the CAG-Cre mice generated Mendelian ratios of WT, fl/+, and CAG-Cre neonates, but the ZBTB38 Δfl/+ was not identified (Table1). Likewise, out of 96 mice born from the fl-neo/+ and CAG-Cre intercrosses, ZBTB38 Δfl-neo/+ neonates were not found, whereas the remaining genotypes displayed the expected Mendelian ratios (Appendix Table S1). These results demonstrate that the loss of a single ZBTB38 allele leads to embryonic lethality.

### Heterozygous loss of ZBTB38 leads to early post-implantation defects

To determine the timing of embryonic lethality, we examined the embryonic morphology at different gestation stages. In an analysis of the littermates from the ZBTB38 fl/+ and CAG-Cre intercross, the expected Mendelian ratios of the four expected genotypes were found to be E6.5-E8.5 (Table1). In contrast, beyond E9.5, no Δfl/+ embryos were identified, whereas the other genotypes were morphologically normal with the expected Mendelian ratio (Table1). Likewise, Δfl-neo/+ embryos were not found beyond E9.5, whereas the other genotypes displayed the expected Mendelian frequency (Appendix Table S1). These results indicated that ZBTB38^+/-^ embryos died around E9.5. Approximately, 70% (n = 9 of 13) of the Δfl/+ embryos were smaller in size than their littermate controls at E6.5 (Fig 3A). At E7.5, the size difference between all Δfl/+ embryos (n = 24 of 24) and controls was more obvious. At E8.5, the Δfl/+ mice were considerably smaller in size and abnormal in morphology compared with the controls (Fig 3A). To further analyze the ZBTB38 heterozygous phenotype, we examined histological sections of embryos using hematoxylin and eosin staining. Compared with littermate controls at E6.5, Δfl/+ embryos showed similar morphological characteristics; however, their epiblasts and ExE were smaller and less organized (Fig 3B). At E7.5, control embryos increased in size and further progressed to form the amniotic, exocoelomic, and ectoplacental cavities. In comparison, all ZBTB38 Δf/+ embryos displayed compact, severely underdeveloped cavities (Fig 3B). At E8.5, all ZBTB38 mutant embryos were significantly smaller than the other genotypes of embryos and were absent from typical organogenesis, which was usually observed in control embryos. Moreover, the embryos exhibited massive red blood cell infiltration (Fig 3B). Collectively, these results indicate that heterozygous loss of ZBTB38 leads to abnormal embryo development starting from E6.5, progressing at E7.5-E8.5, and leading to embryonic lethality at E9.5.

**Figure 3.**
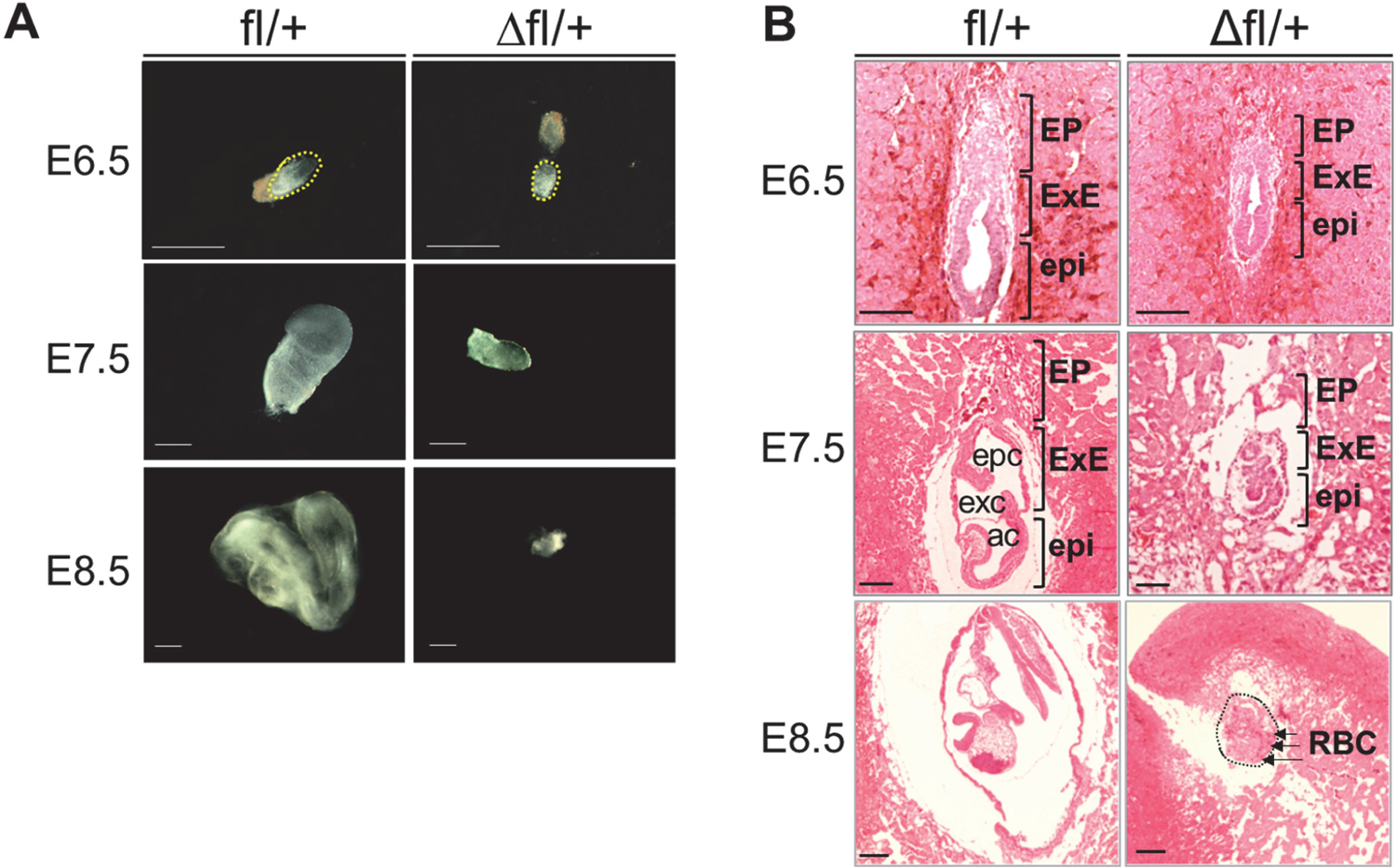
Heterozygous loss of ZBTB38 impairs embryonic development A Morphologies of representative ZBTB38 fl/+ and Δfl/+ embryos at the indicated stages of gestation. Scale bar, 500 mm. B H&E staining for sagittal sections of ZBTB38 Δfl/+ and fl/+ embryos at the indicated stages. Abbreviations: EP: ectoplacental corn, ExE: extraembryonic ectoderm, epi: epiblast, epc: ectoplacental cavity, exc: exocoelomic cavity, ac: amniotic cavity, RBC, red blood cells. Scale Bar: 50 μm.

### Heterozygous loss of ZBTB38 results in decreased proliferation and increased apoptosis in embryos

Because heterozygous mice exhibited a smaller body size, we reasoned that cell proliferation was affected in these embryos. To investigate cell proliferation in the embryos, we performed BrdU incorporation and immunohistochemical analysis of Ki67 in embryos prepared with paraffin sections. In these assays, the cells in the S phase are labeled with BrdU, whereas the cells in the G1, S, G2, and M phases are detected with Ki67. As observed in Fig 4A and B, cells of the control embryos displayed considerable BrdU incorporation in both the epiblast and ExE at E6.5-E8.5 (>80%). In contrast, in the ZBTB38^+/-^ embryo, BrdU-positive cells of epiblast and ExE were dramatically reduced at E6.5 (dropped by 65%), E7.5 (dropped by 57%), and E8.5 (dropped by 34%). Notably, the decrease in BrdU incorporation in the ZBTB38^+/-^ embryos was predominantly observed in the epiblast than in the ExE (Fig 4A and B). In comparison, both the ZBTB38 Δf/+ embryos and control embryos exhibited a high percentage of ki67-positive cells at E6.5-E8.5 (>80%), and no significant difference was observed between them (Appendix Fig S5). Consistent with these results, our previous analysis showed that loss of ZBTB38 functions by gene KO or by siRNA-mediated knockdown in the ES cells inhibited BrdU incorporation, impaired G1 to S transition, and therefore, suppressed proliferation (Nishii *et al*, 2012). Taken together, these findings demonstrate that heterozygous loss of ZBTB38 inhibits G1 to S transition of epiblast cells, and consequently, leads to developmental failure of the embryo.

**Figure 4.**
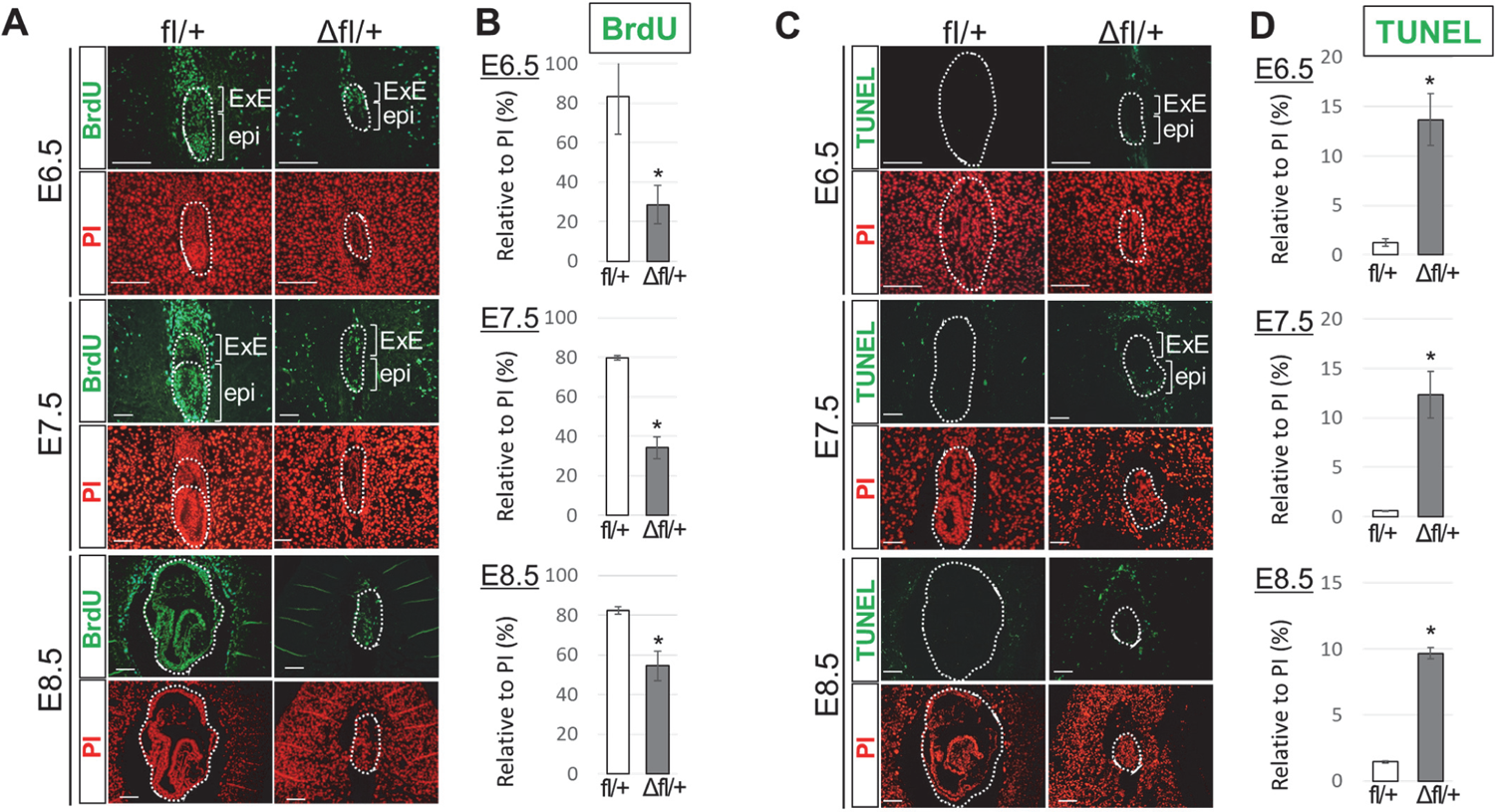
Evaluation of proliferating and apoptotic cells in E6.5-E8.5 embryos A Immunofluorescence analysis of paraffin-embedded sections of controls and ZBTB38 Δfl/+ embryos at E6.5-E8.5. Two consecutive paraffin-embedded sections were taken for performing immunostaining with anti-BrdU antibody (green), and nuclei were counterstained with PI (red). ExE: extraembryonic ectoderm, epi: epiblast. Scale Bar: 50 μm. B Quantitative analysis of the number of labeled BrdU cells relative to the total number of PI-positive nuclei from the indicated umbers of embryos at E6.5 (fl/+: n = 9; Δfl/+: n = 8), E7.5 (fl/+: n = 11; Δfl/+: n = 9), and E8.5 (fl/+: n = 6; Δfl/+: n = 5). Error bars represent ± S.E.M. \**P* < 0.05. C A TUNEL assay was performed on paraffin-embedded sagittal sections from E6.5 embryos onwards. TUNEL-positive cells are shown in green, and nuclei were counterstained with PI (red). Scale Bar: 50 μm. D Quantitative analysis of the number of TUNEL-positive cells relative to the total number of PI-positive nuclei from the indicated umbers of embryos at E6.5 (fl/+: n = 9; Δfl/+: n = 7), E7.5 (fl/+: n = 11; Δfl/+: n = 8), and E8.5 (fl/+: n = 7; Δfl/+: n = 5). Error bars indicate ± S.E.M. \**P* < 0.05.

To explore the fate of the decreased proliferating cells, we speculated that these cells undergo apoptosis. Therefore, we conducted a TUNEL assay, which is the most commonly employed method to detect the fragmented DNA characteristic of apoptotic cells, in paraffin-embedded sections of embryos. As shown in Fig 4C and D, dramatically increased TUNEL-positive nuclei were detected in the ZBTB38 Δf/+ embryos (>10-fold), particularly in the epiblast at E6.5-E7.5, whereas only few such cells were detected in the control embryos at E6.5-E8.5 (<2%). Moreover, cleaved Caspase-3 (cCasp3), another apoptosis marker, was also examined. Similar to the TUNEL analysis, an increased number of cCasp3-positive cells was detected (>10-fold), especially in the ZBTB38^+/-^ epiblast at E6.5-E8.5 (Appendix Fig S6), whereas positive cells were rarely observed in control embryos. Together, these findings indicate that heterozygous loss of ZBTB38 induces apoptosis during early embryonic development and aggravates developmental defects.

### Heterozygous loss of ZBTB38 does not affect the development of 2C to the blastocyst stage but inhibits blastocyst outgrowth

Because ZBTB38 is expressed from the 2-cell to blastocyst stages (Appendix Fig S7), we investigated whether ZBTB38 loss affects pre-implantation development. To test this possibility, 2-cell embryos from the ZBTB38 fl/+ and CAG-Cre intercross were cultured *in vitro* until the 4-cell, 8-cell, and blastocyst stages (Fig 5A and B). Embryos were monitored daily under a microscope, and blastocysts were subsequently analyzed by immunohistochemistry staining and were genotyped. Of the 226 blastocysts assessed, an expected Mendelian ratio was observed among the four genotypes (Fig 5C). During development from the 2-cell to blastocyst stage, the ZBTB38 Δfl/+ embryos appeared to be morphologically indistinguishable from those of their control embryos (Fig 5B). Moreover, proliferation and apoptosis were similar between the Δfl/+ and control blastocysts, as evidenced by the equivalent number of BrdU- and cCasp3-positive cells (Appendix Fig S8A and B). These results suggest that the heterozygous loss of ZBTB38 does not inhibit embryonic development during the pre-implantation stage.

**Figure 5.**
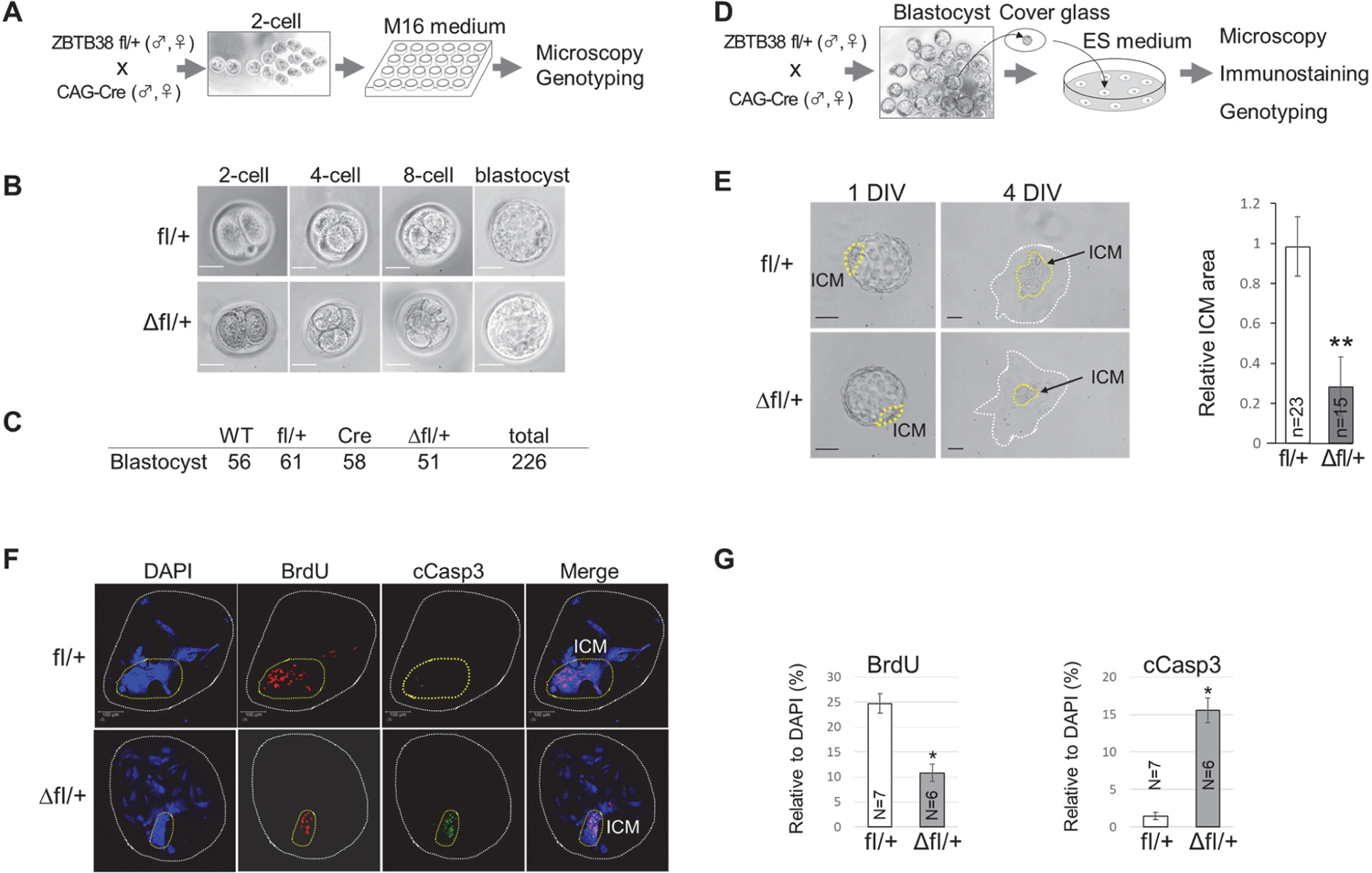
Effects of heterozygous loss of ZBTB38 on pre- and peri-implantation development *in vitro* A Illustration of the 2-cell embryo to the blastocyst *in vitro* culture. B Bright-field microscopy of 2-cell embryo to the blastocyst stage. C Genotypes of blastocysts at day 4 were determined by PCR. D Schematic diagram of embryo outgrowth *in vitro*. E Bright-field microscopy of cultured blastocysts at 1 DIV and 4 DIV are shown (left panel). Yellow dashed lines denote ICM. Scale bar: 100 μm. Right panel, quantitative evaluation of relative ICM area (ICM/TE area). Images were analyze using ImageJ software. The data were representative of five independent experiments and error bars indicate ± S.D. \*\**P* < 0.01. F Confocal immunofluorescence images of BrdU-labeled and cCasp3-positive cells of 4 DIV embryos. DNA was counterstained with DAPI. Yellow dashed lines denote ICM. Scale bar, 100 μm. G Quantitative analysis of the number of BrdU-positive cells (left) or cCasp3-positive cells (right) relative to the total number of nuclei (DAPI-positive cells) from the indicated numbers of representative embryos. Error bars represent ± S.D. \**P* < 0.05.

We next investigated whether loss of ZBTB38 affects blastocyst outgrowth, which recapitulates peri-implantation development *in vivo*. For this purpose, blastocysts obtained from the ZBTB38 fl/+ and CAG-Cre intercross were isolated and cultured for 4 days *in vitro* (DIV) to support further embryonic development beyond the blastocyst stage (Fig 5D). Both ZBTB38 Δfl/+ and control blastocysts normally hatched from zonae pellucidae, attached to the culture dish and initiated outgrowth at 2 DIV, and TE cells differentiated into largely polypoid trophoblast giant cells at 3-4 DIV with no visible difference (Fig 5E and F). Notably, although cells from the control ICM proliferated to expand the area, cells from the Δfl/+ ICM showed a dramatic proliferation defect with a markedly smaller proliferative zone (Fig 5E). To explore whether proliferation and apoptosis account for the aforementioned phenotypes, four DIV embryos were immunostained and analyzed. The results showed that outgrowth of the control blastocysts displayed high proliferation and a low degree of apoptosis, particularly in ICM-derived cells, as evidenced by the high BrdU incorporation with rare cCasp3-positive cells in ICM (Fig 5F). In contrast, a decreased number of proliferating cells and an increased number of apoptotic cells were observed in ZBTB38 Δfl/+ ICM. These data demonstrate that ZBTB38 expression is essential for peri-implantation development.

### Changes in gene expression in ZBTB38^+/-^ embryos and ZBTB38^+/-^ ES cells

To investigate the molecular mechanism by which ZBTB38 loss leads to defective embryos, we focused on E7.5 embryos, which is the stage at which all ZBTB38^+/-^ embryos were dramatically smaller than their controls but could be entirely isolated (Table1 and Fig 3). For this purpose, we carefully isolated the E7.5 fetuses from the intercrosses of ZBTB38 fl/+ and CAG-Cre mice (Fig 6A). Whole-mount immunofluorescence analysis showed that ZBTB38, Nanog, and Sox2, albeit to a lesser extent, were downregulated in the ZBTB38 Δfl/+ embryos as compared to their control littermates (Fig 6B). Moreover, qRT-PCR analysis showed that heterozygotic loss of ZBTB38 reduced ZBTB38 (dropped by 50%), Nanog (dropped by 64%), and Sox2 (dropped by 40%) levels as compared with the controls (Fig 6C). mRNA levels of cyclin E2 and Bcl2, markers of G1/S transition and anti-apoptosis, respectively, were downregulated in the ZBTB38 Δfl/+ embryos as compared to their controls (Fig 6C). Furthermore, the ExE markers of Gata4 and Gata6 were downregulated, whereas the mesodermal gene, Brachyury, was not affected. Collectively, these findings demonstrate that loss of ZBTB38 single allele results in a half decrease of ZBTB38 *in vivo*, giving rise to misregulation of pluripotency, proliferation, differentiation, and apoptotic genes.

**Figure 6.**
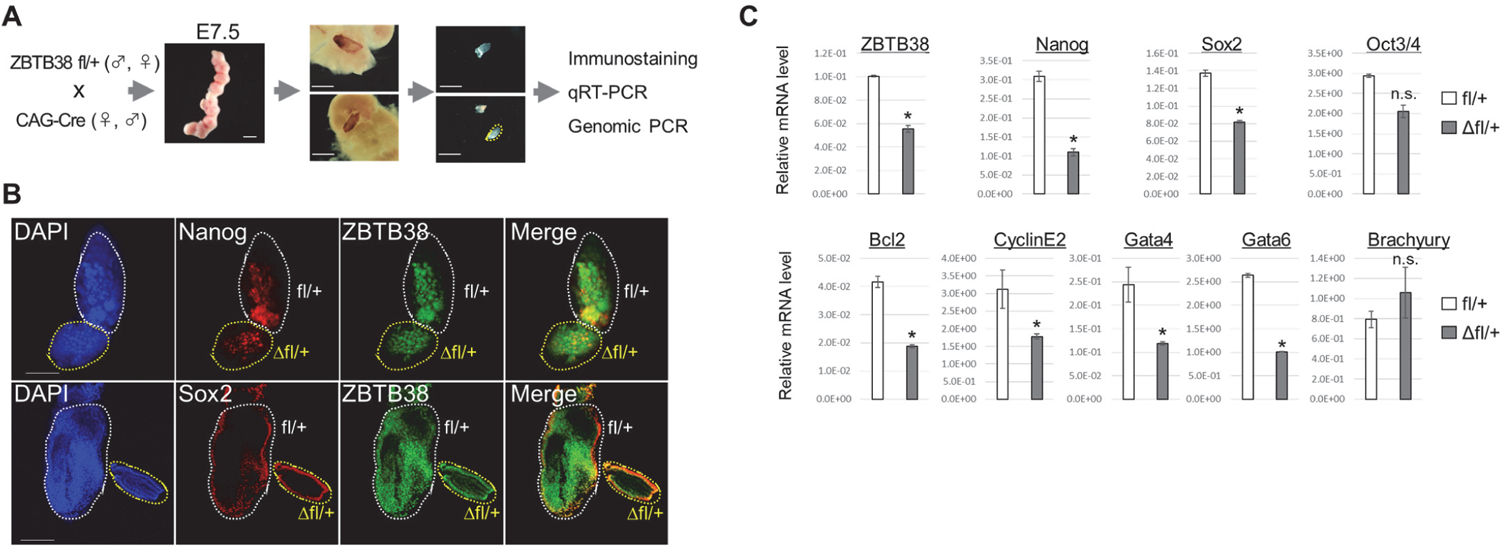
Expression levels of ZBTB38 and markers in E7.5 embryos A Schematic diagram of isolated E7.5 fetuses. B Confocal images of whole-mount immunohistochemical staining. Expressions of the indicated proteins were detected by anti-ZBTB38, anti-Nanog, and anti-Sox2 antibodies. Cell nuclei were counterstained with DAPI. Scale bar denotes 50 μm. **C** qRT-PCR results of the indicated gene expressions of the ZBTB38 Δfl/+ embryo and control embryo. Data shown are representative of three independent experiments and error bars represent ± S.D. \**P* < 0.05.

To investigate the effects of ZBTB38 loss on ES cell pluripotency and self-renewal, blastocysts from the ZBTB38 fl/+ and CAG-Cre intercross were cultured in ES medium to generate ES cells (Fig 7A). ES colonies derived from WT, ZBTB38 fl/+, and Δfl/+ blastocyst cultures were established without differences in frequencies (data not shown), indicating that ZBTB38 is dispensable for the generation of ES cells. We found that the ZBTB38^+/-^ ES cells retained their morphological features of undifferentiated ES cells, stained similarly positive for alkaline phosphatase activity, indicative of the pluripotent, undifferentiated state (Fig 7B). To further characterize these ES cell properties, the expression of pluripotent genes was examined by western blotting and qRT-PCR. As shown in Fig 7C and D, ZBTB38 mRNA and protein levels in the Δfl/+ ES cells were reduced to half as compared to those of their controls, indicating that the loss of a single ZBTB38 allele led to a half decrease in its expression *in vitro*. Moreover, the mRNA and protein levels of Nanog and Sox2, except those of Oct4, were downregulated in the ZBTB38^+/-^ ES cells as compared to those in the controls (Fig 7C and D).

**Figure 7.**
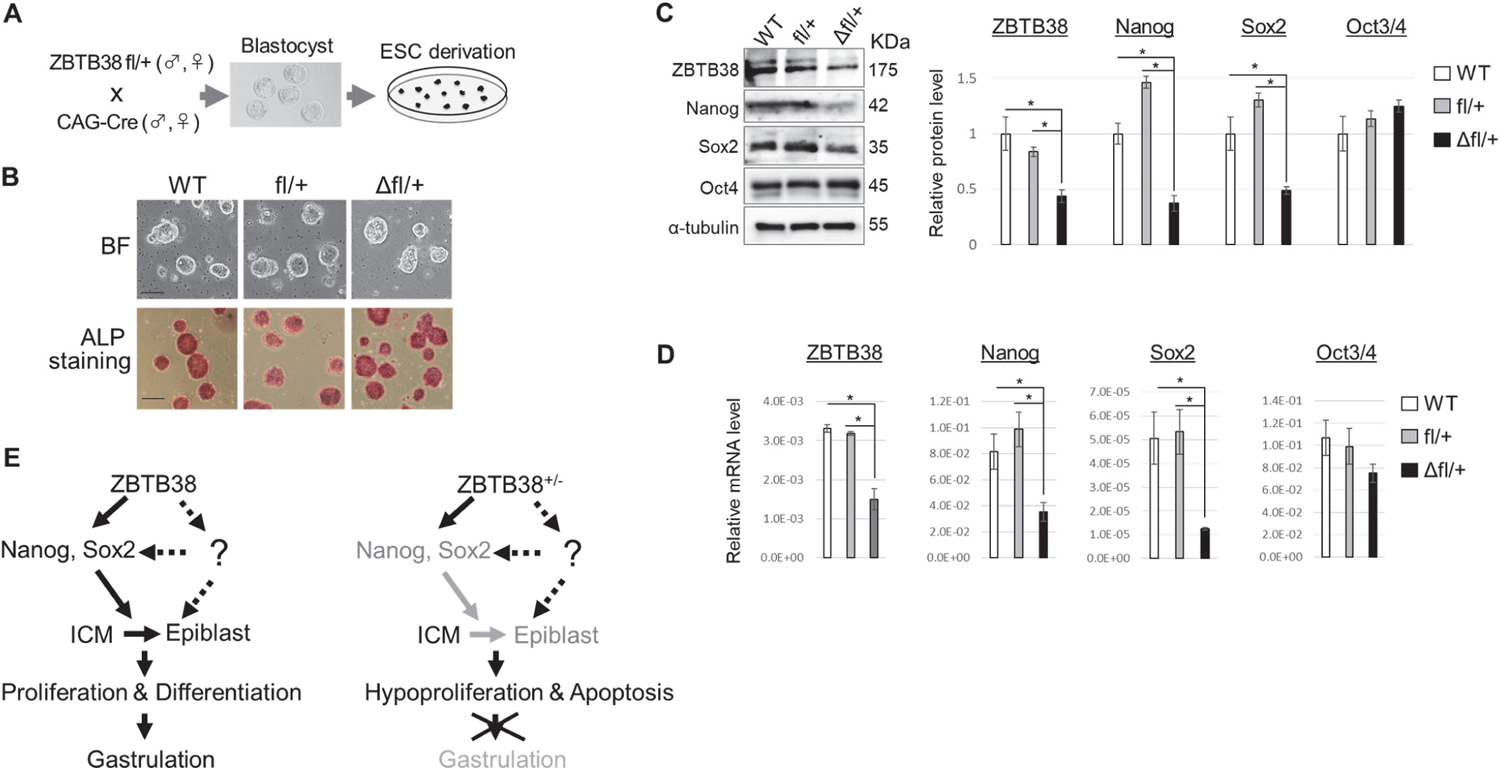
Establishment of ES cell clones and expressions of genes in ES cells A Schematic diagram showing derivation of ESCs from blastocysts. B Bright-field images of typical ES cell colonies (top) which show compact colonies with distinct borders and well-defined edges, and alkaline phosphatase staining (bottom) of ESCs. Scale bar, 100 μm. C, D Expressions of the indicated mRNA (C) and protein (D) from ES cells. *GAPDH* and α-tubulin were used as internal controls for qRT-PCR and immunoblotting, respectively. Data shown are representative of three independent experiments, and error bars indicate ± S.D. \**P*, < 0.05. **e** Schematic model of how ZBTB38 affects gastrulation.

Altogether, these findings demonstrate that heterozygous loss of ZBTB38 did not affect pre-implantation development, but instead suppressed ICM outgrowth during peri-implantation. Heterozygous loss of ZBTB38 results in the downregulation of Nanog and Sox2, and this decrease leads to hypo-proliferation and apoptosis in the epiblasts, resulting in gastrulation defects (Fig 7E).

## DISCUSSION

Our results from the two different strategies demonstrated that loss of the ZBTB38 single allele led to embryonic developmental failure. In agreement with the developmental phenotypes of Dnmts KO mice, our results showed that ZBTB38^+/-^ embryos also died around E9.5. We presented the data indicating that loss of ZBTB38 decreased the expression of genes responsible for epiblast proliferation, differentiation, and cell viability.

Currently, the physiological necessity of MBP during peri-implantation remains almost unknown despite MBPs being expressed at varying levels throughout embryonic stages (Buck-Koehntop & Defossez, 2013; Martin Caballero *et al*, 2009). Here, we provide genetic evidence that ZBTB38 is required for early embryogenesis. Because of the complexity of methylation pattern dynamics and technical limitations at the peri-implantation stage, where, when, and how MBP binds to the methyl-CpG sites of genes, transposons, and genome, remain unknown. Ubiquitin-like with PHD and ring finger domains 1 (Uhrf1, also known as Np95) specifically binds to hemi-methylated CpG sites and recruits DNMT1 to these sites to inherit DNA methylation patterns during DNA replication. Similar to Dnmt1^-/-^ mice, Uhrf1^-/-^ embryos died shortly after gastrulation (Sharif *et al*, 2007). The following findings demonstrate that ZBTB38 is unlikely to have the same mechanism as Uhrf1. (1) Uhrf1 binds to hemi-methylated CpG and Dnmt1, whereas ZBTB38 binds to a single symmetrical methylated CpG (CGCCAT or GCGGTA) (Sasai *et al*, 2010) and currently lacks direct evidence of ZBTB38 binding to Dnmt1. (2) Both Uhrf1^-/-^ and Dnmt1^-/-^ ES cells exhibited genome-wide demethylation (Sharif *et al*, 2007), whereas the ZBTB38^-/-^ ES cells did not (unpublished data), suggesting that ZBTB38 is dispensable for establishing or maintaining DNA methylation in ES cells. Further experiments using methyl-ATAC-seq or/and methyl-ChIP-seq are required to explore the ZBTB38 binding dynamics and properties in ES cells and early embryos (WT vs. heterozygote). In addition, the DNA methylation-dependent requirement of ZBTB38 in the established methylation sites of genes, transposons, and genome during peri-implantation remains unknown. Generation of knock-in mutant mice in which ZBTB38 is unable to bind to methylated CpG is expected to answer this question.

To our knowledge, only two of the known 6014 gene KO databases (https://www.mousephenotype.org/), the vascular endothelial growth factor (VEGF) and the Notch ligand DLL4, have been shown to exhibit heterozygous embryonic lethality. VEGF^+/-^ and DLL4^+/-^ mice, generated by a conventional targeting strategy, died around E10.5 because of vascular developmental failure (Ferrara *et al*, 1996; Gale *et al*, 2004). In this study, both the conventional and Cre-loxP-based conditional knockout approaches revealed that heterozygous loss of ZBTB38 resulted in embryonic lethality. These findings provide new insights for understanding of heterozygous embryonic lethality.

Wong et al. generated ZBTB38^-/-^ mice using CRISPR/Cas9 technology in a C57BL/6 background (Wong & Bhattacharya, 2020). The ZBTB38 fl/fl mice were crossed with CMV-Cre mice expressing Cre recombinase ubiquitously under the control of a human cytomegalovirus promoter to obtain germline ZBTB38 deletion. Unexpectedly, ZBTB38^-/-^ mice were born, developed normally, and were fertile, with no detectable abnormalities of massive gene expression. However, expression levels of either the mRNA or protein of ZBTB38 was not demonstrated in their KO mice, implying that the normal phenotype is unlikely to be the consequence of lacking ZBTB38.

The results presented in this study indicate that ZBTB38 gene haploinsufficiency downregulated Nanog and Sox2 expression. It has been shown that either the Nanog^-/-^ or Sox2^-/-^ mice died soon after implantation, whereas their heterozygous mice appeared normally and were fertile(Mitsui *et al*, 2003; Avilion *et al*, 2003), suggesting that their dosage is critical for embryogenesis. Based on the following findings, we speculated that the loss of Nanog and Sox2 expression accounts, at least partially, for the phenotypic outcome of ZBTB38^+/-^ embryos. First, ZBTB38 largely colocalized with Nanog and Sox2 in the ICM of blastocysts (Fig 1B), and partially colocalized with them in the epiblast of E6.5 embryos (Fig 1C). Second, immunofluorescence data showed that Nanog and Sox2 were downregulated in the E7.5 ZBTB38^+/-^ embryos (Fig 6B). Third, qRT-PCR results revealed that heterozygous loss of ZBTB38 led to reduction of Nanog and Sox2 expression in E7.5 embryo (Fig 6C) and in ICM-derived ES cells (Fig 7C and D). Fourth, our previous data have revealed that ZBTB38 positively regulates RF8 ES cell (derived from 129/TerSv mice) proliferation by regulating its downstream target Nanog, and the phosphoinositide 3-kinase (PI3K) signaling pathway accounts for this regulation (Nishii *et al*, 2012). Notably, the loss of ZBTB38 (knockdown, heterozygous, and homozygous KO) in RF8 ES cells decreased the expression of Nanog, but not of Sox2 (Nishii *et al*, 2012). Strain differences in ES cells (129 vs. C57BL6) may explain this discrepancy, although further experiments are required. In this study, we also observed that heterozygous loss of ZBTB38 downregulated the expression of endodermal genes (Gata4 and Gata6), but did not reduce the expression of mesodermal genes at E7.5 (Fig 6C). Gata4/Gata6 is essential for PrE formation and is critical for vascularization and nutrient transport during embryo development (Morrisey *et al*, 1998; Molkentin *et al*, 1997; Cai *et al*, 2008). It is likely that the loss of ZBTB38 impairs PrE cell differentiation to form a functional yolk sac, thus disturbing embryo development.

To date, the molecular mechanism by which ZBTB38 regulates Nanog and Sox2 expression remains unclear. In ES cells, Nanog activates Sox2 transcription, and vice versa. In embryos, however, Nanog is not required to initiate transcription of Sox2, and vice versa. Both the Nanog promoter and the Sox2 regulatory region are largely unmethylated in blastocysts and ESCs, but they are methylated in differentiated cells (Mitsui *et al*, 2003; Okita *et al*, 2007), suggesting that DNA methylation accounts for their expressions. Molecularly, ZBTB38 may activate Nanog and Sox2 transcription by binding to their promoters or regulatory regions in a spatiotemporally defined manner, or by recruiting unknown factor(s), which, in turn, modulates their expression (Fig 7E). Because Nanog promoter contains two ZBTB38 consensus binding sites (the methylated CGCCAT and non-methylated CAGGTG), biochemical experiments are undertaken to explore how ZBTB38 regulates Nanog expression.

ZBTB38 is ubiquitously expressed in tissues (Sasai *et al*, 2005), and its expression is associated with height (Gudbjartsson *et al*, 2008), cancers (Kote-Jarai *et al*, 2011; Jing *et al*, 2019; de Dieuleveult *et al*, 2020; Chen *et al*, 2019) and neurodegenerative diseases (Mead *et al*, 2012; Brown *et al*, 2014). Moreover, down-regulation of ZBTB38 expression potentiates the toxicity of anti-tumor reagents in cancer cells (Marchal *et al*, 2018). Thus, the generation and analysis of the tissue-specific Cre-mediated KO (hetero and homo) mice will comprehend ZBTB38’s physiological functions and ZBTB38-associated diseases.

## MATERIALS AND METHODS

### Generation of ZBTB38 fl/+ ESCs and ZBTB38 △fl/+ mice

Targeting vector was purchased from European Conditional Mouse Mutagenesis Program (EUCOMM, ID: MAE-2331), encompassing a 25-kilobase (kb) DNA fragment including 10 kb of ZBTB38 homologous sequence in which the 5′ and 3′ arms of homology are 6.2 kb and 4.1 kb, respectively. The recombinant allele containing an FRT-flanked LacZ and neo cassettes was linearized with *AsiS*I and was electroporated into 129-derived ES Cells. After electroporation, ES Cell clones were grown in the presence of 500 μg/ml geneticin and isolated after culturing for 8-12 days on mitomycin C-treated SNL-STO cells as described previously (Kotoku *et al*, 2016). The correct integration of the targeting construct and homologous recombination was confirmed by sequencing and restriction enzyme digestion, and genomic PCR (gPCR), respectively. ES cells were cultured in ES medium (DMEM, 15% fetal bovine serum, 2 mM L-glutamine, 100 μM nonessential amino acids, 1% penicillin and streptomycin, and 0.1 mM β-mercaptoethanol) supplemented with 1000 IU/ml recombinant murine leukemia inhibitory factor (LIF, Nacalai Tasque) as described previously (Kotoku *et al*, 2016).

The proper recombinant ES clone was injected into C57BL/6J blastocysts with subsequently transplantation into pseudo-pregnant female mice. Chimeric males were selected based on coat color and subsequently crossed with C57BL/6J female mice to generate the ZBTB38 fl-neo/+ mice. Male fl-neo/+ mice were crossed to the C57BL/6J females to generate F1 offspring. The ZBTB38 fl-neo/+ mice were crossed with FLP mice to remove the FRT-flanked splice acceptor sites to generate ZBTB38 fl/+ mice. The ZBTB38 fl/+ mice were crossed with a CAG-Cre strain that ubiquitously expresses the Cre recombinase to obtain the ZBTB38 Δfl/+ mice. ZBTB38 fl/+ mice were backcrossed with C57BL/6J at least for eight generation.

A single male was paired with one or two females, which were plug checked and weighed daily. The embryonic days were counted starting E0.5 on the day the vaginal plug was detected. C57BL/6J mice were purchased from CLEA (Japan). All mice were approved by the Animal Care Committee of Nara Institute of Science and Technology and conducted in accordance with guidelines that were established by the Science Council of Japan.

### Genotyping

The genomic DNA was prepared from the ear punching with Gentra Puregene Cell Kit (Qiagen), or from embryos at various developmental stages with Gentra Puregene Cell Kit or AllPrep DNA/RNA FFPE Kit (Qiagen). All experiments were carried out according to the manufacturer’s instructions. Approximately 1-100 ng of DNA was used for genotyping by PCR analysis. Information of primers was shown in Appendix table S3.

### 2-cell and blastocyst collection and ES cell establishment

Isolation and establishment of blastocyst was performed according to the protocol described (Bryja *et al*, 2006) with slight modification. Briefly, 2-cell or blastocyst was obtained from natural mating or superovulation. For superovulation, female mice (4-8 weeks old) were intraperitoneal injected with 7.5 IU pregnant mare’s serum gonadotrophin (PMSG), followed 48 h later by an intraperitoneal injection of 7.5 IU of human chorionic gonadotrophin (hCG), both were purchased from ASKA Pharmaceutical Co., Ltd. Female mice were mated with males immediately after hCG injection. Approximately 44 h after hCG injection, 2-cell embryos were flushed from oviduct at E1.5 and cultured in M16 medium (M7292, Sigma-Aldrich) covered with mineral oil until they reached blastocyst stage. For blastocysts isolation, E3.5 embryos were flushed from uterine horns, and cultured with M16 medium under a 5% CO2, 37°C incubator.

For generating ES cells, individual late-cavitating blastocysts of similar morphology from each experimental group were treated with Tyrode’s solution (Sigma-Aldrich) to remove zona pellucida, then transferred onto the SNL-STO feeder cells. Blastocysts were cultured in ES medium with 1000 IU/ml LIF and the differentiation inhibitors 3i (3 μM CHIR99021, 1 μM PD0325901 and 10 μM SB431542, all were purchased from Selleck). After five days of culture, ICM outgrowths were individually collected, dissociated with trypsin-EDTA, and transfer the cells to 24-wells plated on feeder layers. The resulting undifferentiated colonies were further propagated in the same medium. Putative ES cell lines were passaged several times in the ES cell medium, genotyped, characterized by microscopy.

### Blastocyst outgrowth assay

The blastocyst outgrowth assay was performed as described (Soares *et al*, 2013), with slight modifications. Briefly, blastocysts with or without zona pellucida were plated into a 0.1% gelatin-treated 24-well plates and cultured in ES medium lacking LIF. Embryos was imaged daily with NIKON microscope. On day 4 (4 DIV), embryos were fixed with 4% paraformaldehyde, immunostained and imaged, then the DNA was extracted and genotyped by PCR. Areas from the 4 DIV ICM and TE outgrowths were calculated from images using ImageJ software, and the ICM ratio (defined as ICM area/TE area) was calculated using Excel.

### Alkaline phosphatase assay

Alkaline phosphatase (AP) activity of undifferentiated ES cell was detected using the Alkaline Phosphatase Staining Kit II (STEMGENT), according to manufacturer’s instructions.

### Embryo dissection and histological analysis

Morphological and histological analysis of embryos were collected at various times of gestation. For histological analysis, embryos were rinsed with ice-cold PBS and fixed with 4% paraformaldehyde at room temperature for 10-30 min, dehydrated through graded alcohols and embedded in paraffin. Embryos were sectioned at 5 µm thickness and stained with hematoxylin and eosin. Sectioned embryos were photographed using a NIKON microscope. For immunofluorescent analysis, paraffin sections were rehydrated, immunostained with cleaved Caspase-3 antibody (#9664, Cell Signaling), Ki-67 antibody (#12202, Cell Signaling). Antigen retrieval was performed according to the individual antibody instructions. CF488A Donkey anti-Rabbit IgG (#20015, Biotium) and CF488A Donkey anti-Mouse IgG (#20014, Biotium) were used as the secondary antibody. Nuclei were counterstained with 1 μg/ml Propidium Iodide (P4170, Sigma-Aldrich), washed and mounted.

Terminal deoxynucleotidyl transferase (TdT) mediated dUTP nick end labeling (TUNEL) assay was performed according to the manufacturer’s instructions (MBL, #8445). Briefly, paraffin-embedded sections were deparaffinized, DNA nick end labelled, and counterstained with 1 μg/ml Propidium Iodide and mounted. The slides were analyzed using a TCS SP8 confocal microscope (Leica).

### BrdU incorporation assay

For 5-bromo-2-deoxyuridine (BrdU; Sigma-Aldrich) incorporation, pregnant female mice at specific times were injected intraperitoneally with BrdU (50 μg/g of body weight). The mice were sacrificed by cervical dislocation after 2 hrs injection. For histological analysis, embryos and tissues were fixed in 4% paraformaldehyde overnight at 4°C and embedded in paraffin wax for sectioning. Five-micrometer sections were deparaffinized, and incubated with anti-BrdU antibody (#66241-1, Proteintech) after heat antigen retrieval and denaturation according to the manufacturer’s instructions. Anti-Mouse CF488A was used as the secondary antibody. After staining, the slides were counterstained with 1 μg/ml propidium iodide. Blastocysts or blastocysts from outgrowth were treated with 20 μM BrdU for 2-4 hrs before paraformaldehyde fixed and immunostained.

### Whole-Mount Immunohistochemistry for embryos

Whole-Mount Immunohistochemistry was performed as described (Joyner & Wall, 2008) with a slight modification. Briefly, embryos (blastocysts, blastocyst from outgrowth, E6.5-E7.5 embryos) were washed three times with cold-PBS, and fixed with methanol/DMSO (4:1) at 4° C overnight. Embryos were treated with methanol/DMSO/H_2_O_2_ for 3 hrs, and rehydrated with 75%, 50% and 25% methanol. After 30 min of incubation with 0.3% Triton X-100 in PBS, embryos were blocked with the washing buffer (0.2% Tween 20 in PBS) supplemented with 10% goat serum at room temperature for 120 min or overnight at 4° C. After washing several times with the washing buffer, embryos were incubated with anti-ZBTB38 antibody (Oikawa *et al*, 2011), anti-Nanog (#560259, BD Phamingen), anti-Sox2 (S1451, Sigma-Aldrich), anti-Oct4 (sc-5279, SantaCruz) for 120 min, washed with washing buffer, and then incubated with secondary antibody for 60 min. Secondary antibodies used were Alexa Fluor 594 goat anti-Mouse IgG (#8890, Cell Signaling) and Alexa Fluor 488 goat anti-Rabbit IgG (A32731, Thermo Fisher). Nuclei were counterstained with 4′,6-diamidino-2-phenylindole (DAPI). After washed with washing buffer, signal was viewed under a confocal laser-scanning microscope (TCS SP8, Leica). All procedures were performed in a six-well plate except for the experiment with outgrowth embryos, which was performed in a four-well plate.

### RNA extraction and quantitative Real-Time PCR

Total RNA was extracted using Sepasol (Nacalai Tasque) for beyond E9.5 embryo, AllPrep DNA/RNA FFPE Kit (Qiagen) for paraffin section, PicoPure™ RNA Isolation Kit (Thermo Fisher) for blastocyst-E8.5 embryos, according to the manufacturer’s instructions. cDNA was synthesized using ReverTra Ace® qPCR RT Master Mix with gDNA Remover (TOYOBO). Quantitative real-time PCR (qRT-PCR) was performed using a LightCycler® 96 System (Roche Diagnostics) with the Thunderbird SYBR Green PCR Mix (Toyobo), following the manufacturer’s instructions as described^32^. 2-μl aliquots of all cDNA samples were analyzed in triplicate on 96-well optical PCR plates (Roche Diagnostics). GAPDH or TBP was used as the reference gene and all analyses were performed using the ΔΔCt method with Roche LightCycler 96 system software. Primer sequence are listed in the Table S2.

### Western Blotting

Western blotting was performed as described previously (Sasai *et al*, 2005). Briefly, protein lysates were prepared in RIPA buffer supplemented with complete protease inhibitor (Roche Applied Science). Proteins were then separated on 6%–15% SDS-PAGE, transferred onto PVDF membranes, and probed with anti-ZBTB38 (Oikawa *et al*, 2011), anti-Oct3/4 (MAB1759, R&D Systems), anti-Sox2 (S1451, Sigma-Aldrich), anti-Nanog (AB5731, Millipore), anti-α-tubulin (clone DM 1A, Sigma-Aldrich) antibodies. HRP-conjugated anti-mouse or anti-rabbit IgG (Cell Signaling) were used as secondary antibodies. Quantification was conducted by using GelQuant.NET software provided by biochemlabsolutions.com.

## Statistical analysis

Unless stated otherwise, data are given as means ± standard deviation (SD). Each experiment included at least three independent samples and was repeated at least three times. Statistical analyses were performed with GraphPad Prism 7.0 software using two-tailed unpaired Student’s *t*-test, and differences were considered significant when \**p* < 0.05, \*\**p* < 0.005. “n.s.” indicates not significance (*p*>0.05).

## Data availability

No data were deposited in public database.

## ACKNOWLEDGMENT

We are grateful to Drs. Masashi. Kawaichi, Yasumasa. Bessho, Hiroshi. Ito and Ayako. Isotani for valuable suggestions and discussion. We would like to thank Messrs. Anton and Fumiyuki Mukaite, Mrs. Eri Hosoyamada for technical assistance, and the members of the Ishida laboratory for suggestions and technical advice. This work was supported by a Grant-in-Aid for Scientific Research (16K08587) from the Japan Society for the Promotion of Science (JSPS).

## Conflict of Interests

The authors declare no conflict of interest exists.

## CONTRIBUTIONS

M.N. and T.M. were involved in the conception and design of the study, performed the experiments, data analyses and collected and assembled data. S.H. and S.O. conducted the experiments; C.O., N.S. and Y.I. offered assistance of the animal experiments, technical guidance and advices; E.M. was involved in the conception, design, performed the experiments, data analyses, and was responsible for manuscript writing and final approval of the manuscript. All authors read and approved the manuscript.

---

**Figure S1.**
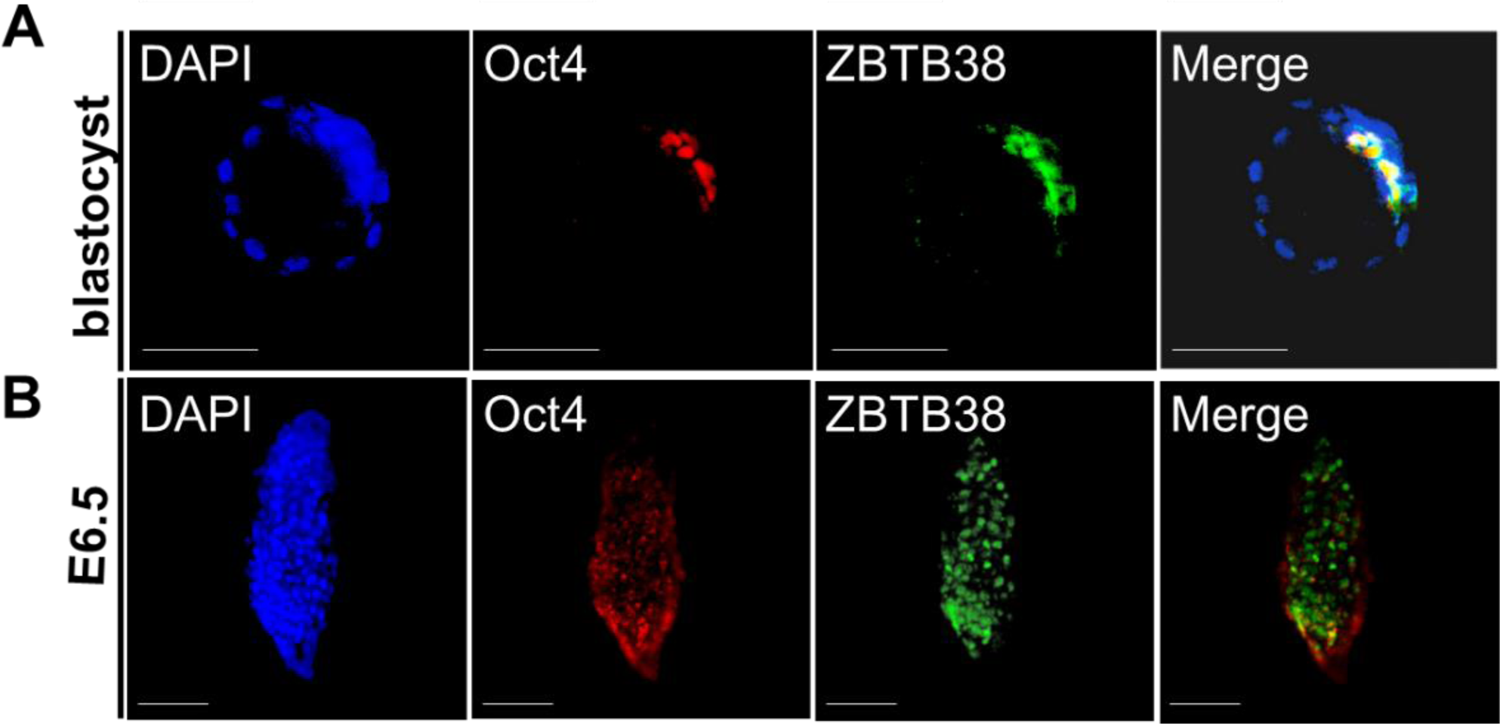
Expression patterns of ZBTB38 **A** Whole-mount immunofluorescence and confocal microscopy for blastocyst (**A**) and E6.5 (**B**). Expression patterns of the indicated proteins were detected by anti-ZBTB38 and anti-Oct4 antibodies. Cell nuclei were counterstained with DAPI. Scale bar denotes 50 μm (1A) or 100 μm (1C). ExE: extraembryonic ectoderm, epi: epiblast.

**Figure S2.**
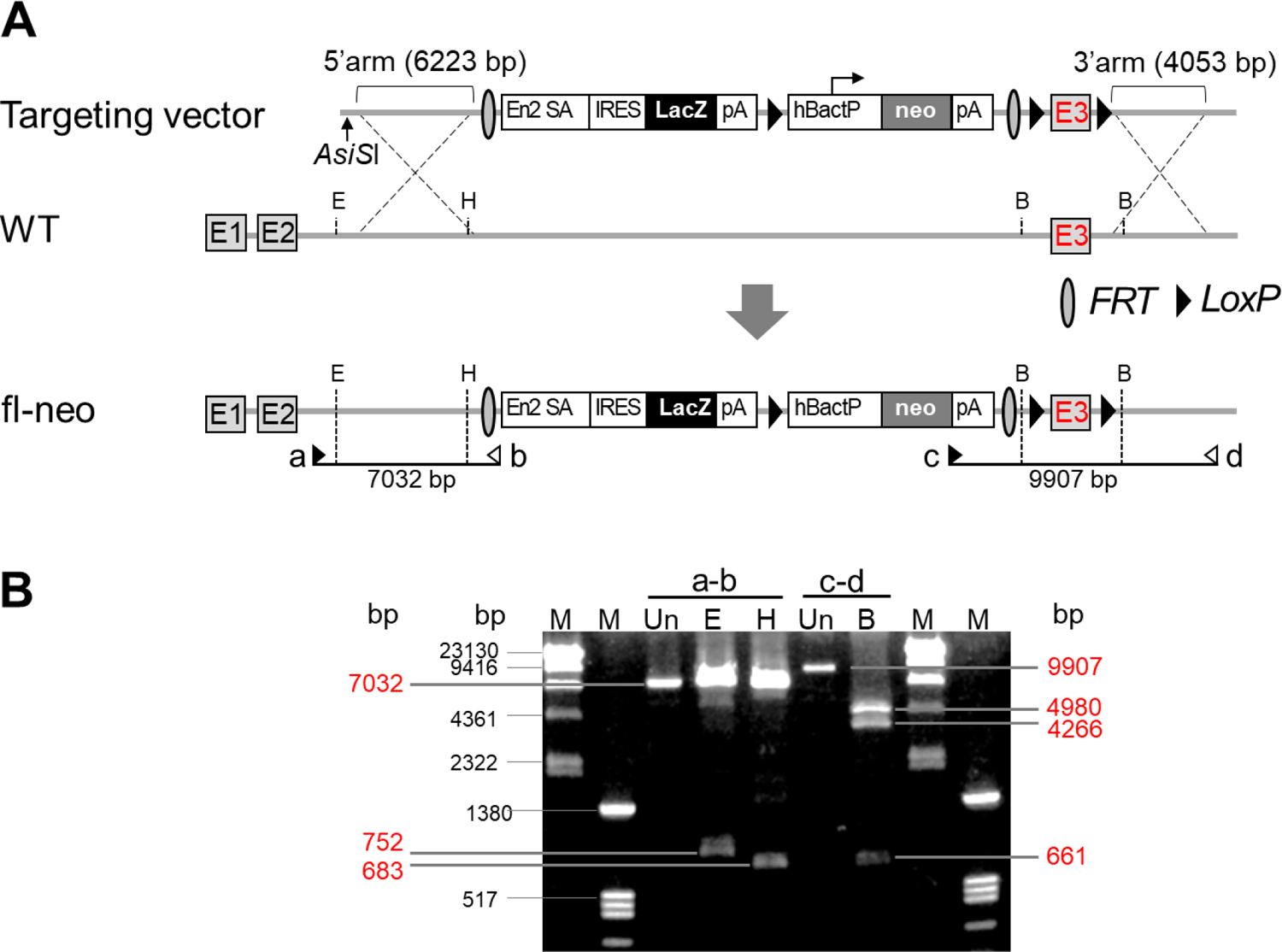
Schematic representation of the targeting vector and locus A Schematic of EUCOMM targeting vector. The vector contains 10kb of ZBTB38 homologous sequence in which the 5′ and 3′ arms of homology are 6.2 kb and 4.1 kb, respectively. Homologous recombination in ES cells results in the generation of the fl-neo targeted allele. Abbreviations, SA: splice acceptor, IRES: internal ribosome entry site, pA: poly(A) signal, En2 SA: mouse En2 splicing acceptor, IRES: internal ribosome entry site, hBactP: human b-actin promoter. Primers (a-d) are shown by arrows and the expected size (bp) of genomic PCR are shown under the individual primer pairs. B Results of genomic PCR for confirmation of homologous recombination. Red number denotes the expected size of bands digested with the indicated restriction enzymes. E: *EcoR*I, B: *BamH*I, H:*Hind*III, Un: uncut. M denotes size marker.

**Figure S3.**
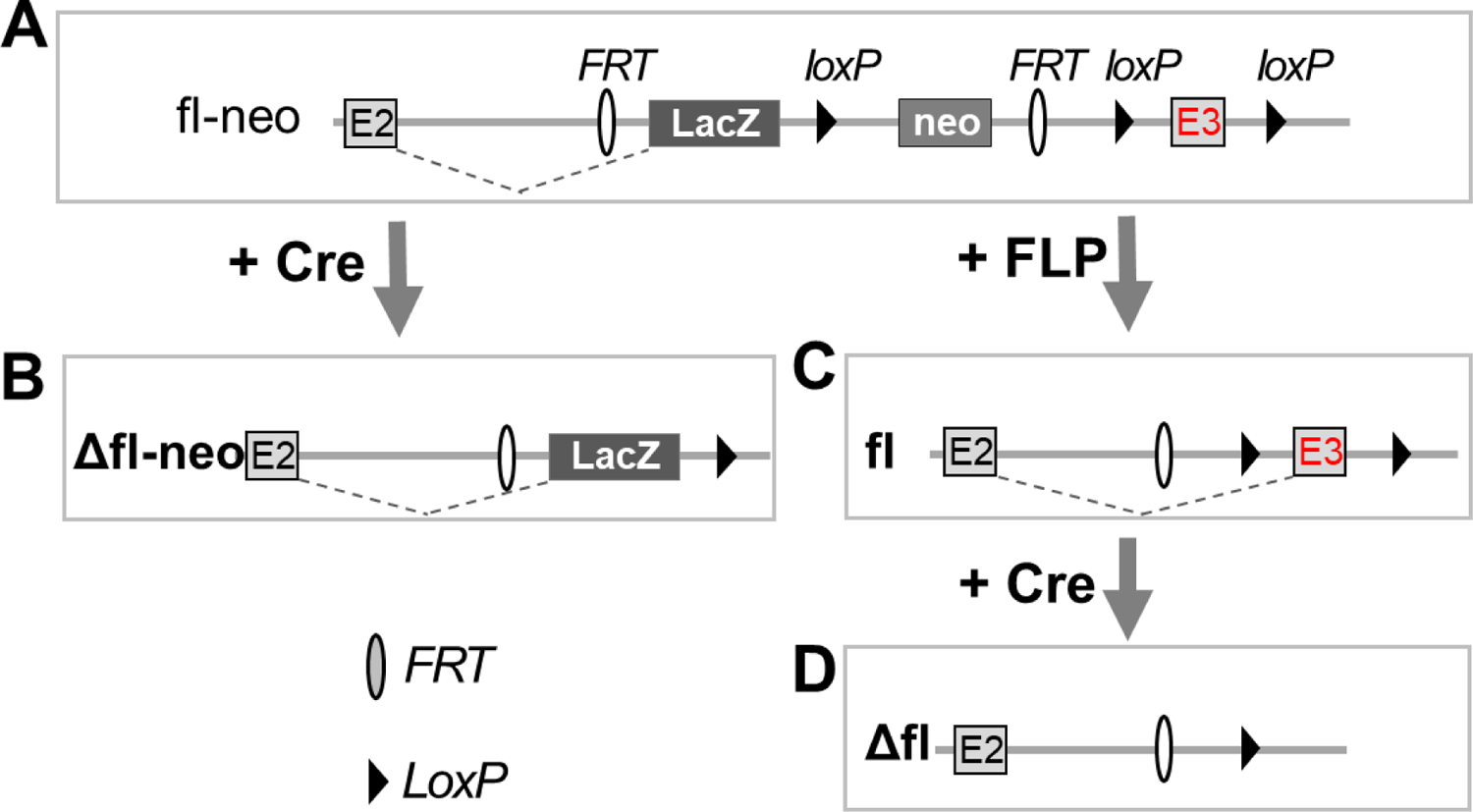
Two strategies to generate ZBTB38^+/-^ allele A fl-neo targeted allele. B Crossing the fl-neo allele to a mouse strain expressing CAG-Cre recombinase, yields a lacZ-tagged allele lacking ZBTB38 exon 3 and neomycin cassette (Δfl-neo). C Crossing the fl-neo allele to a mouse line expressing FLP recombinase results in a conditional-ready allele lacking both the lacZ and neomycin cassettes (fl). D Crossing the fl allele a mouse strain expressing CAG-Cre recombinase leads to a null allele lacking ZBTB38 exon 3 (Δfl). Exons are shown as empty boxes and marked by a number inside. Neo: neomycin-resistant gene; FLP: flp recombinase.

**Figure S4.**
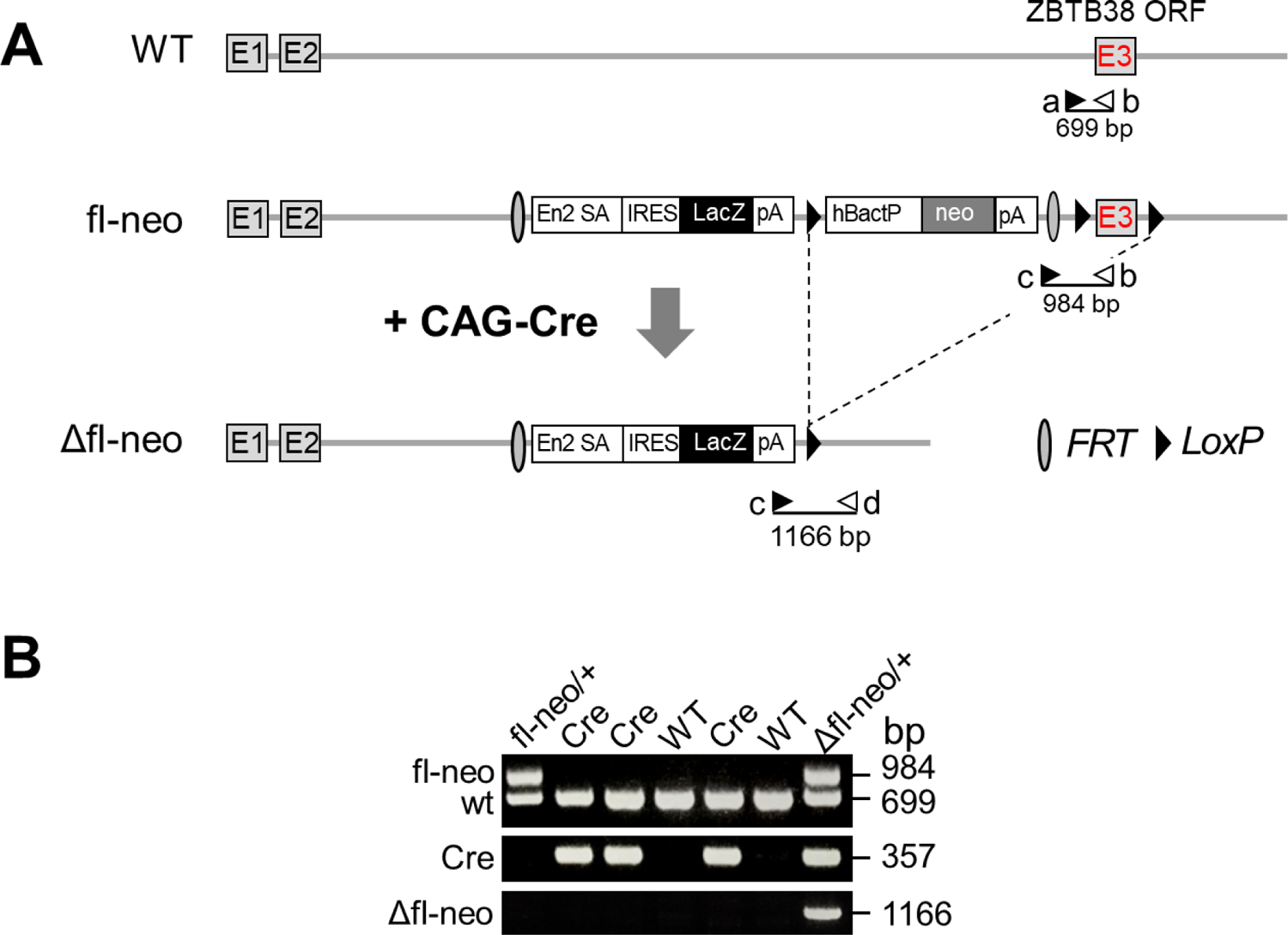
Generation of the ZBTB38 Δfl-neo/+ mice A Schematic representation of WT and knockout constructs. The deleted allele (Δfl-neo) lacking the ZBTB38 exon 3 is produced by CAG-Cre-induced recombination as shown. The position of the primers (a∼d) used for genotyping is shown with arrows, the expected size (bp) of genomic PCR are shown under the individual primer pairs. B PCR genotyping of E7.5 embryos isolated from the ZBTB38 fl-neo/+ mice and CAG-Cre mice intercrosses. WT, fl and Δfl alleles produced a 699-bp, 984-bp and 1166-bp band, respectively. Cre genotyping primer produces a 357 bp band.

**Figure S5.**
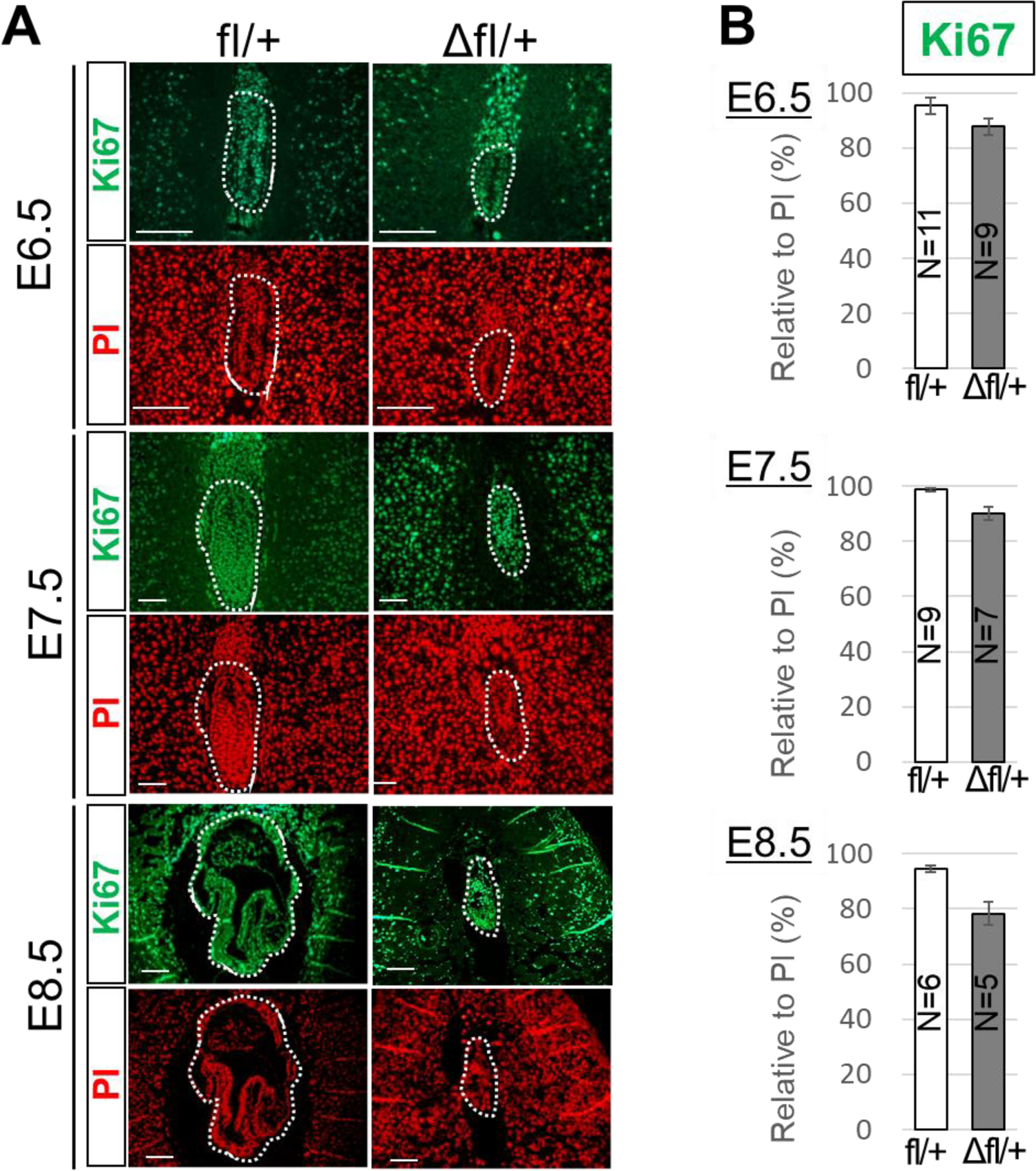
Evaluation of proliferating cells in embryos A The number of Ki67-positive cells in paraffin-embedded sagittal sections of E6.5-E8.5 embryos. Immunofluorescence analysis of control ZBTB38 fl/+ (a1∼a6) and ZBTB38 Δfl/+ (b1∼b6) embryos. Paraffin-embedded sections were taken for performing immunostaining with anti-Ki67 antibody (green) and nuclei were counterstained with PI (red). Scale Bar: 50 μm. B Quantitative analysis of the number of labeled Ki67 cells relative to total number of nuclei (PI-positive cells) from the indicated umbers of representative sections. Error bars represent ± S.E.M.

**Figure S6.**
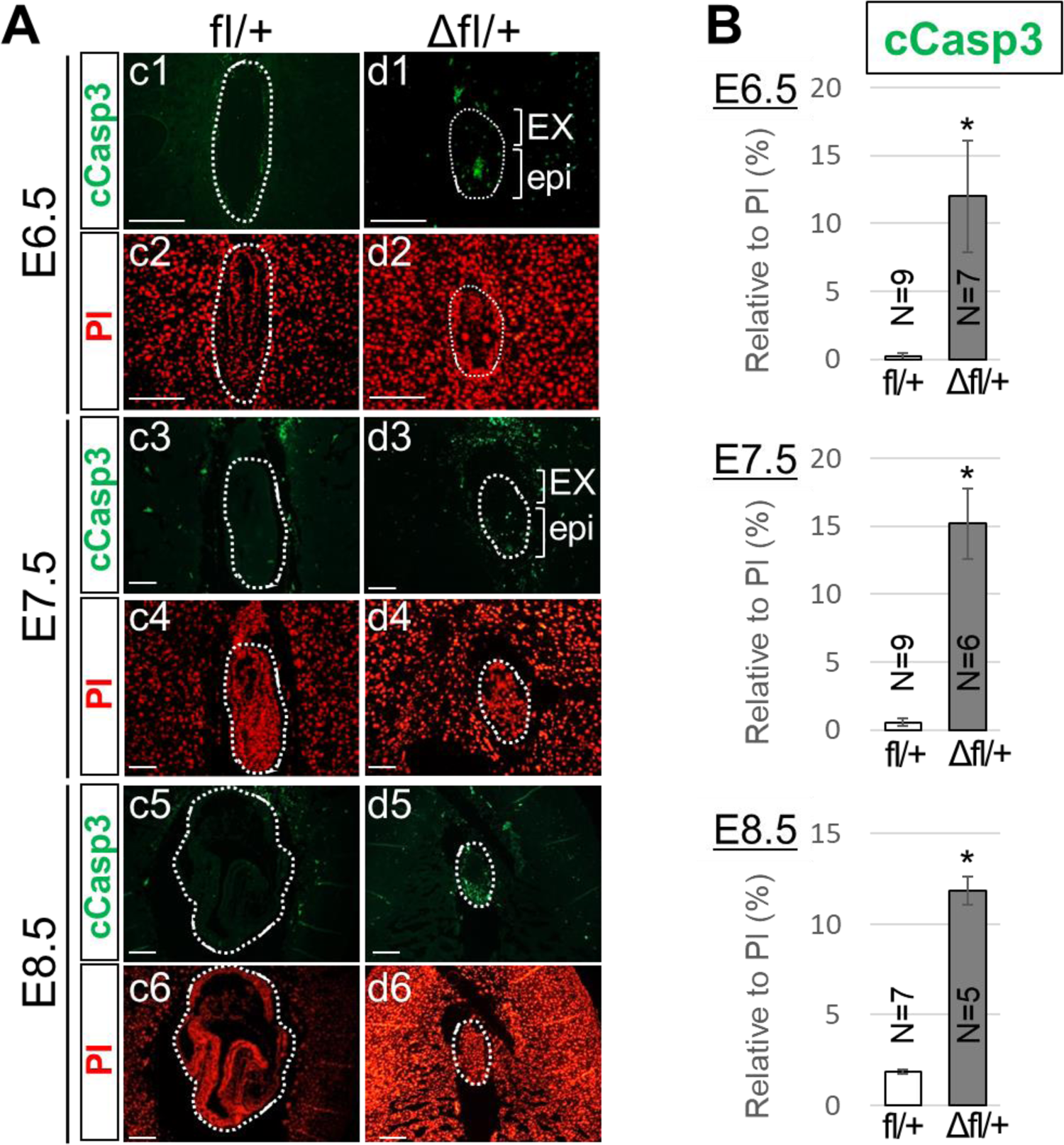
Evaluation of apoptotic cells in embryos A Immunofluorescence assay was performed on paraffin-embedded sagittal sections from E6.5 onwards. cCasp3-positive cells are shown in green and nuclei were counterstained with PI (red). Scale Bar: 50 μm. B Quantitative analysis of the number of cCasp3-positive cells relative to total number of nuclei (PI-positive cells) from the indicated umbers of representative sections. Error bars indicate ± S.E.M. \**P* < 0.05

**Figure S7.**
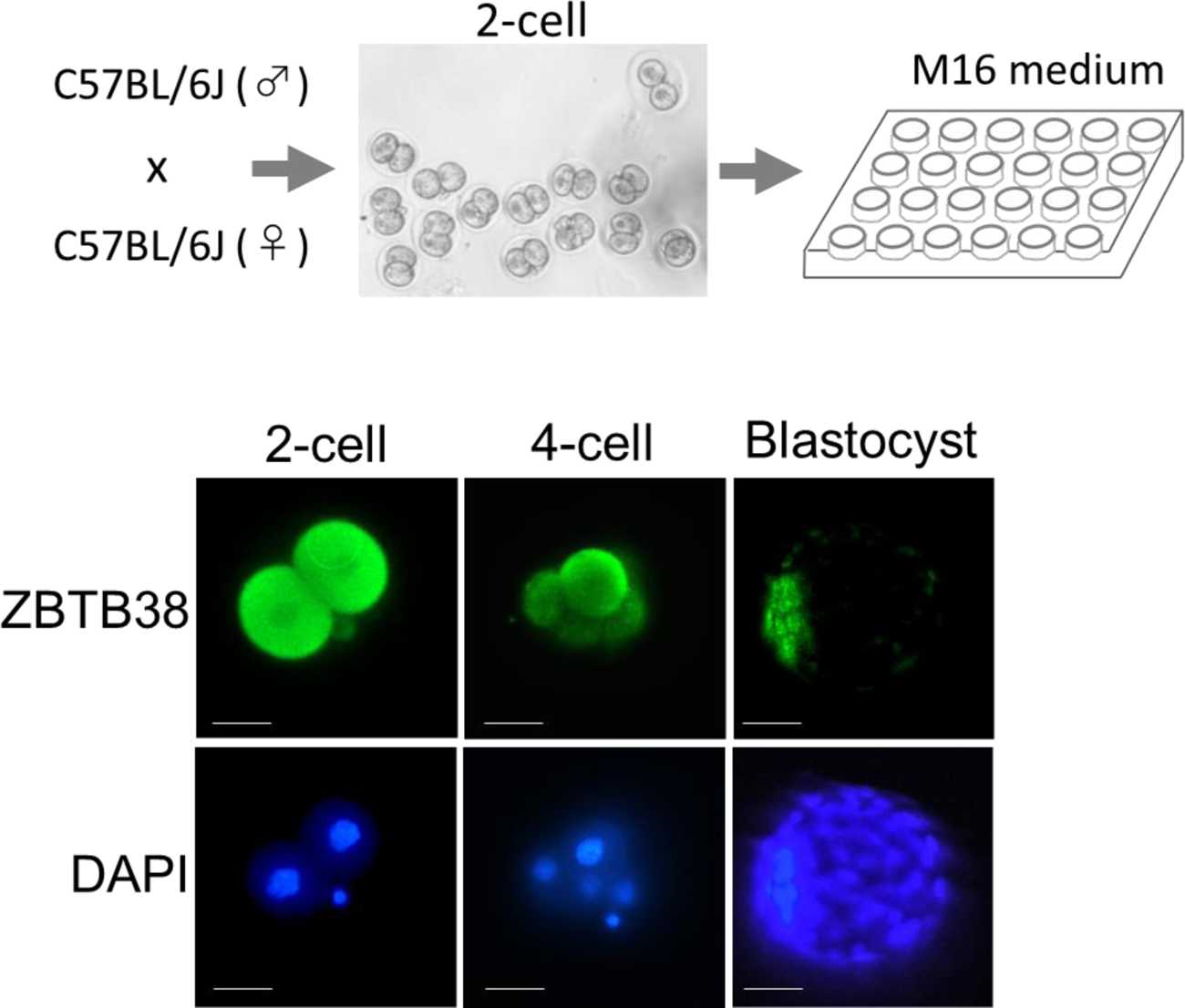
Expression patterns of ZBTB38 in pre-implantation embryos Illustration of culturing 2-cell embryos to blastocysts *in vitro* (upper panel). Lower panel, immunofluorescence microscopy for ZBTB38 expression with anti-ZBTB38 antibody at the indicated stage of embryos. Cell nuclei were counterstained with DAPI. Scale bar denotes 50 μm.

**Figure S8.**
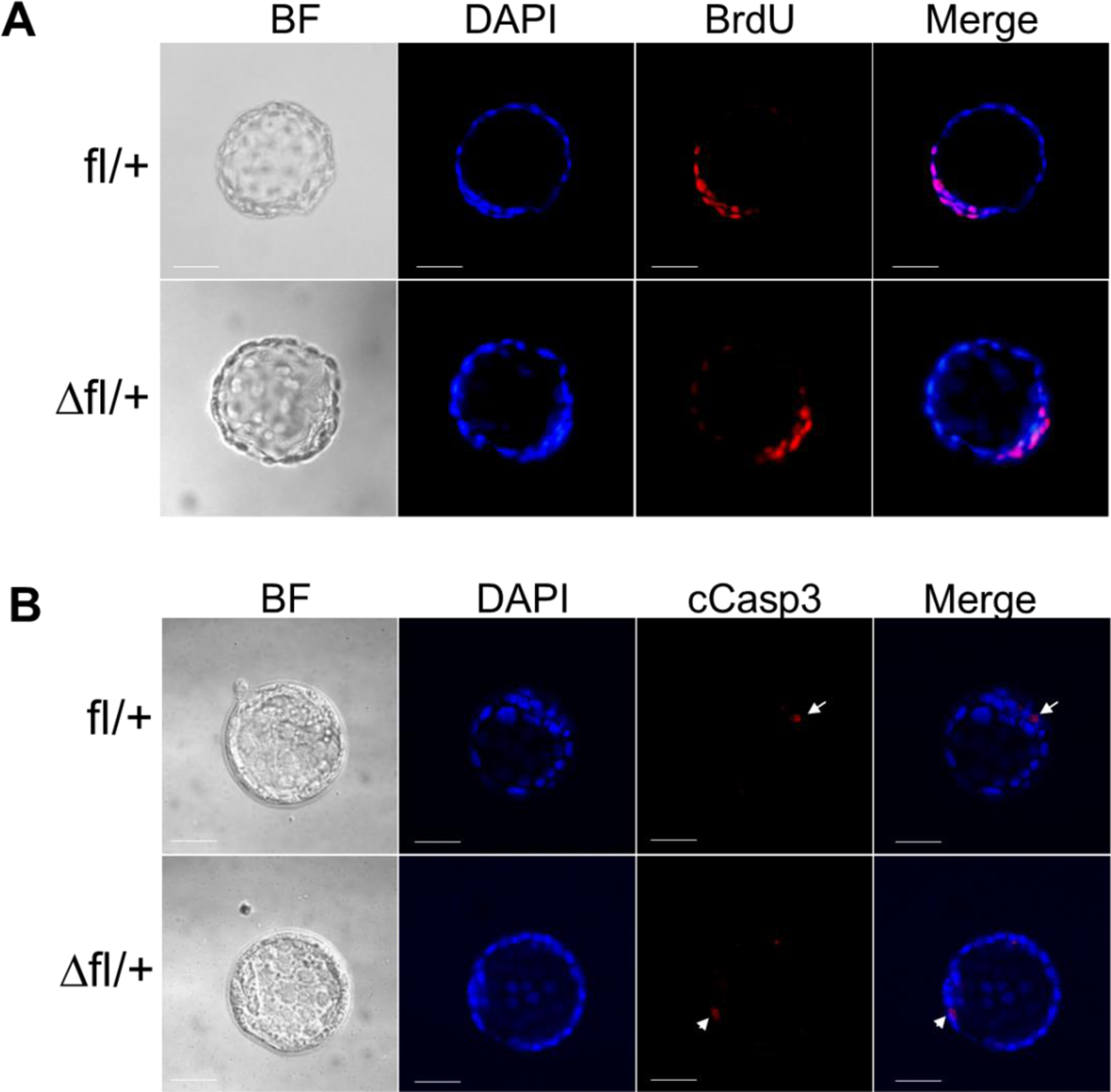
Evaluation of proliferation and apoptosis in blastocyst A, B Whole-mount immunofluorescence and confocal microscopy of blastocysts from the ZBTB38 fl/+ mice and CAG-Cre mice intercrosses. Immunostaining with anti-BrdU antibody (red, A) and anti-cCasp3 antibody (red, B) are shown. Arrows indicate cCasp3-positive cells. Nuclei were counterstained with DAPI (green). Scale bar, 50 μm.

**Appendix Table S1.** Genotype analysis of offspring from the ZBTB38 fl-neo/+ and CAG-Cre intercrosses

| **stage** | **Total** | **WT** | **fl-neo/+** | **Cre** | **Δfl-neo/+** |
| --- | --- | --- | --- | --- | --- |
| **Newborn** | **96** | **37** | **30** | **29** | **0** |
| **E10.5** | **35** | **12** | **12** | **11** | **0** |
| **E9.5** | **30** | **6** | **14** | **7** | **3 (#3)** |
| **E8.5** | **30** | **9** | **4** | **10** | **7 (\*7)** |

#absorbed; \* abnormal

**Appendix Table S2.**
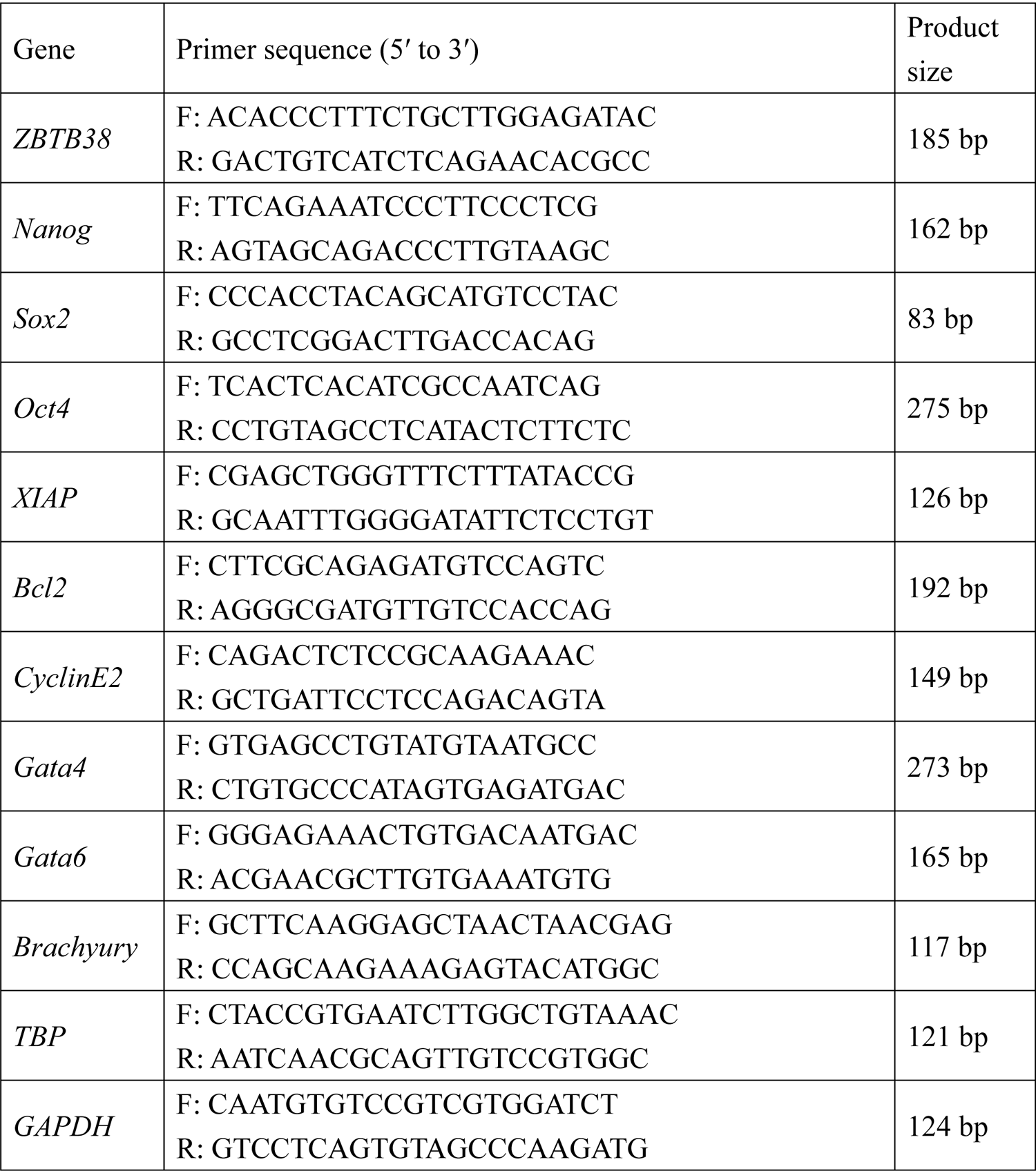
Primer used for qRT-PCR

**Appendix Table S3.**
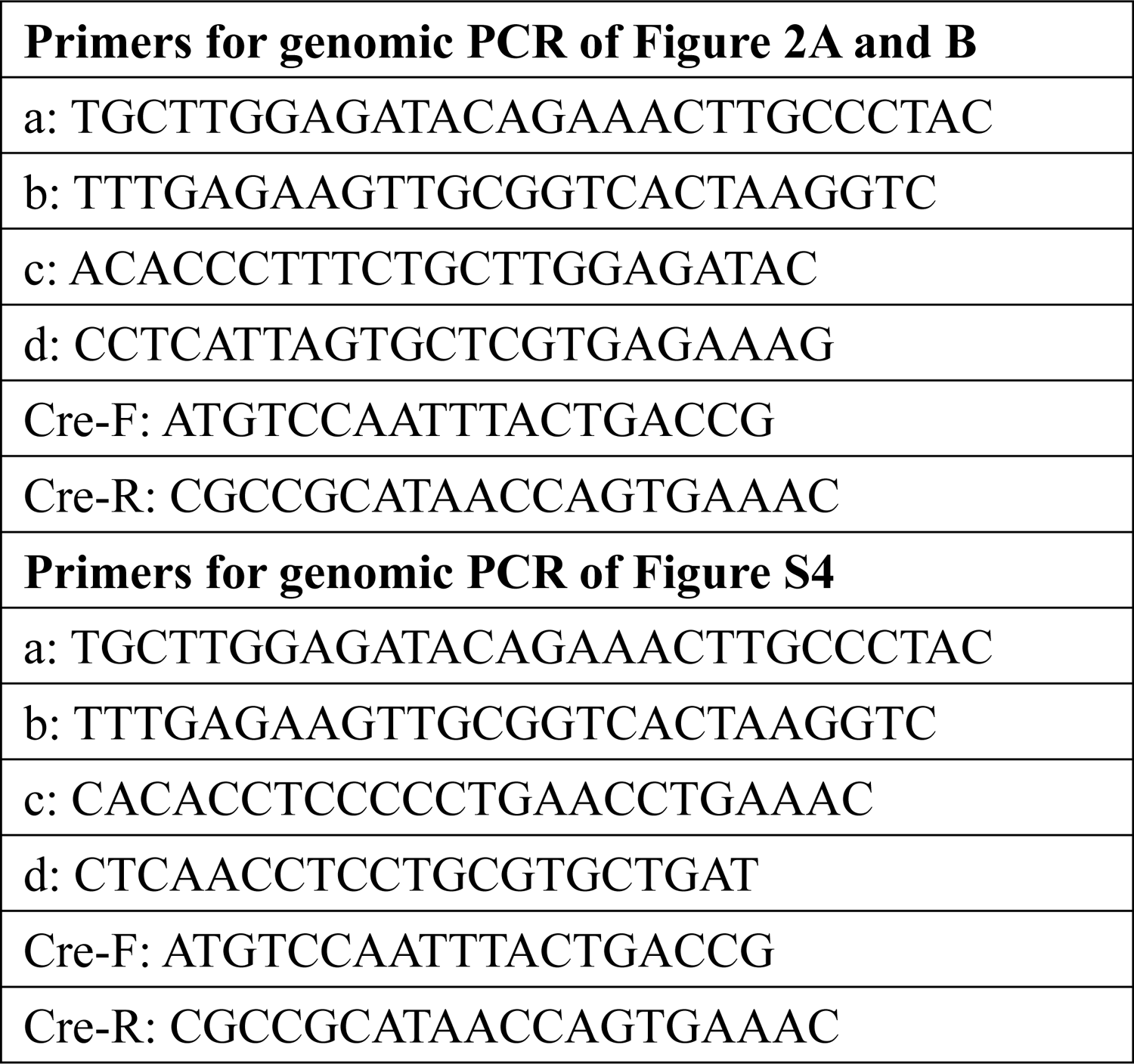

